# Enriched experience increases reciprocal synaptic connectivity and coding sparsity in higher-order cortex

**DOI:** 10.1101/2025.09.01.673156

**Authors:** Rajat Saxena, Justin L. Shobe, Aida M. Andujo, Wing Ning, Christelle Anaclet, Bruce L. McNaughton

## Abstract

The integration of new information during sleep reshapes cortical representations that support categorical knowledge. Auto-associative attractor network theories predict that reciprocal excitatory connections help form stable categorical attractors, but direct evidence is missing. We tested this using ten weeks of enriched experience (ENR) in mice as a model for knowledge accumulation and recorded single-unit activity across hippocampus and neocortex. ENR induced significant remodeling in high- but not low-level neocortex, with a shift from unidirectional to bidirectional excitatory-excitatory connections, suggestive of increased ‘cell assemblies’. This was accompanied by increased inhibitory-to-excitatory connections and sparser, more orthogonal population activity during awake rest and slow-wave sleep, particularly in deep layers. Thus, ENR reorganizes cortical circuits into a symmetric, inhibition-balanced network that improves coding efficiency, supporting long-standing attractor network predictions.

## Introduction

In the neocortex (NC), new knowledge is integrated with prior information, which is organized into abstract cognitive schemas or concepts. These schemas enable the flexible recombination of previously learned representations, which facilitates the rapid integration of new information (i.e. forward knowledge transfer). However, the network mechanisms underlying schema storage and organization remain poorly understood.

Hebbian cell assembly theory and modern auto-associative attractor network models, including Hopfield networks, predict that increasing the number of stored patterns leads to more reciprocal (bidirectional) synaptic connectivity (*1–4*). Such connectivity enhances robustness to noise, enables signal amplification, and supports pattern completion (*5*). These models also predict that increasing representational sparsity expands memory capacity by reducing overlap between stored patterns – trends consistently observed in artificial neural networks, where both sparsity and orthogonality increase across learning epochs (*6*).

Here, we tested the hypothesis that environmental enrichment (ENR) (*7–9*), as a model of knowledge accumulation, would produce neural changes consistent with the foregoing theoretical predictions. Specifically, we predicted that knowledge-rich brains would exhibit increased bidirectional excitatory connectivity, compared to exercise-only control brains. We used a recently developed ENR protocol where animals are exposed to complex, multisensory, and stimulating conditions that enhance both behavioral performance (*10*) and neural functions (*11–13*) and combined it with high-density electrophysiology recordings across the retrosplenial cortex, primary visual cortex, and hippocampus (HC). This approach allowed us to examine how enriched experiences alter functional synaptic connectivity and population coding dynamics, particularly during offline periods such as awake rest and slow-wave sleep (SWS), when the brain freely explores its internal state-space (*14–17*).

### Experimental Paradigm

We ran n=18 (2-month-old, 10M/8F) pair-housed mice on either an enrichment track (ET; n=5M/4F) or an exercise control track (CT; n=5M/4F) for 10-12 weeks (5 x 1-hour sessions per week) (Fig. 1A-C). ET mice were exposed to multiple obstacles that were changed daily to allow a rich repertoire of visual, somatosensory, and motor experiences, while CT mice ran on a track with simple ramp hurdles (20 cm x 11 cm per hurdle) (Fig. S1A, B). CT mice ran more laps than ET mice (F=5.938, p=0.033, group-effect, repeated-measures ANOVA (rmANOVA)) (Fig. 1C). A separate group of n=15 mice (7ET/8CT, age-matched to main group) underwent the same ENR protocol and confirmed that ET group showed enhanced spatial recognition in a Y-maze task (t=−4.2, p=0.0013, unpaired t-test; Fig. S1C-F), replicating previously shown behavioral benefits of ENR using this paradigm (*10*).

**Fig. 1.**
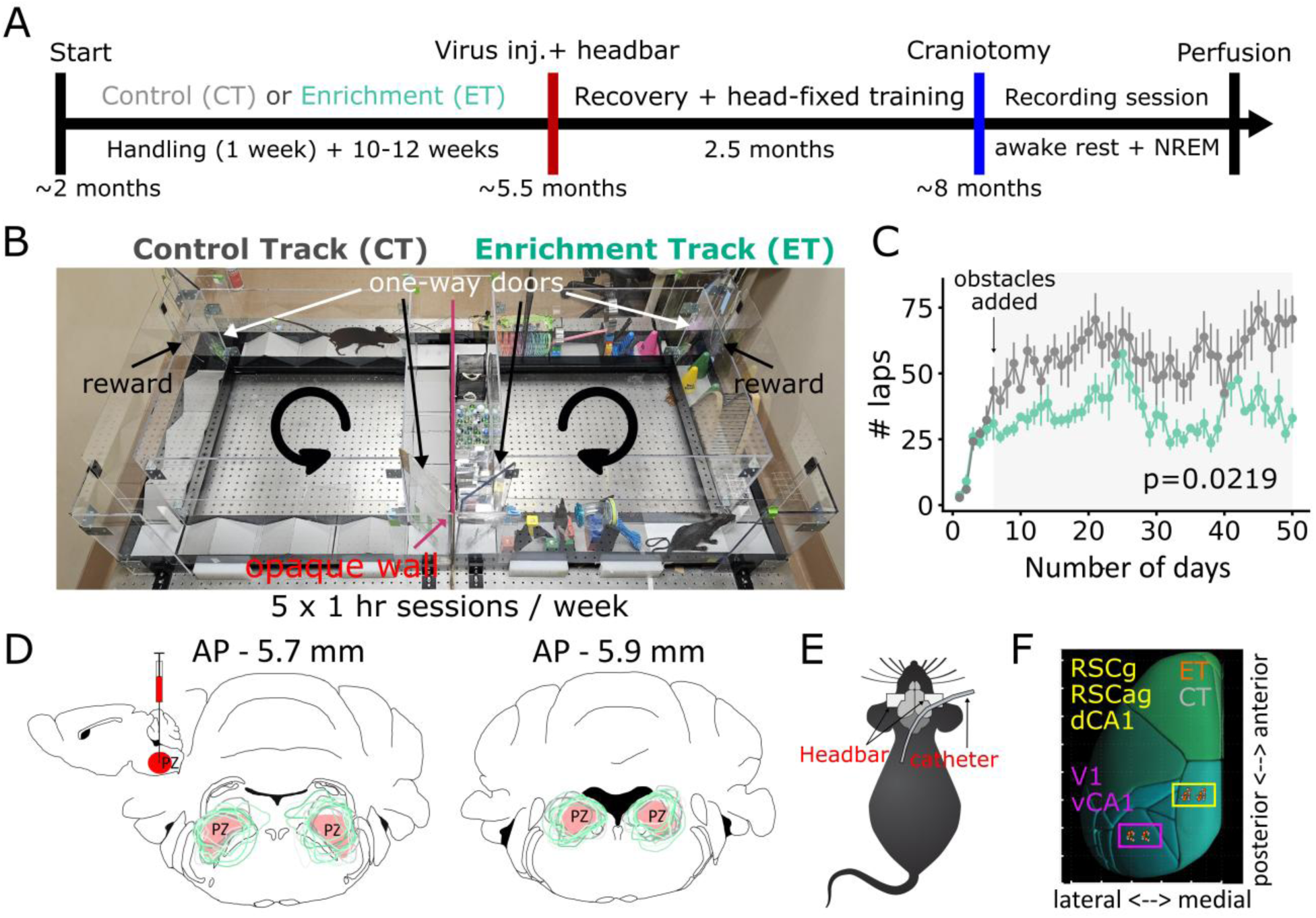
Environmental enrichment (EE) protocol and experimental design. (**A**) Timeline: ∼2-month-old mice ran for 10-12 weeks on either an enrichment (ET, aquamarine) or control track (CT, gray) (track: 86.5cm long x 86.5cm wide; arm width: 11.43cm). After enrichment/control treatment, animals were head-barred and injected with DREADDs into the Parafacial Zone (PZ) to enable chemogenetic induction of SWS. After ∼2-2.5 months of head-fixation training, craniotomies were performed over RSC, V1, and CA1, and a catheter was implanted for subcutaneous CNO delivery. Each mouse underwent one recording session: 2.5-3 hours of awake rest followed by ≥ 13 hours of induced SWS. (**B**) We ran double-housed young adult mice (8 weeks) in our ENR setup such that one mouse ran on CT (left), while the paired cagemate ran on ET (right) for 5 x 1-hour sessions/ week for 10-12 weeks. ET mice encountered 12 obstacles (new obstacle configuration every session), while CT mice ran over the same simple ramp hurdles. Arrows indicate running direction. (**C**) Mean number of laps ± sem for 10 weeks (50 days). ET mice ran fewer laps than CT (F=5.938, p=0.033, rmANOVA). (**D**) Inset: schematic showing pAAV2-hSyn-DIO-hM3D(Gq)-mCherry injected (bilaterally) into the PZ (red). Virus expression for both ET (green) and CT mice (gray) for two coronal sections. Schematic of (**E**) catheter implant and (**F**) dual-probe craniotomies: dCA1, DG, RSCag, RSCg (yellow box); V1, vCA1 (pink box). Colored circles (ET=orange and CT=gray) mark each animal’s recording sites for both probes (mirrored for recording from other hemisphere).

After ENR, mice (∼5.5 months old) were head-barred and injected bilaterally with pAAV2-hSyn-DIO-hM3D(Gq)-mCherry virus in the parafacial zone (PZ), a SWS promoting region, to induce essentially natural SWS at will using Clozapine-N-oxide (CNO) (*18*) (Fig. 1D). This allowed stable data collection for long-duration offline periods, which is required for reliable estimation of connections between excitatory neurons (*19*). After ∼2 months of head-fixation habituation, craniotomies were performed over granular and agranular retrosplenial cortex (RSCg, RSCag), primary visual cortex (V1), dentate gyrus (DG), and dorsal and ventral CA1 (dCA1, vCA1), and a catheter was implanted in the upper back for subcutaneous CNO injection (Fig. 1E-F).

Acute electrophysiology recordings were conducted the day after craniotomy surgery using two dual-shank 256-channel silicon probes (512 total) (*20*), targeting RSC-dCA1 (probe 1) and V1-vCA1 (probe 2). Recordings included 2.5-3 hours of awake rest, followed by at least 13 hours of CNO-induced slow-wave sleep (SWS), with CNO boosters administered subcutaneously (0.5 mg/kg), every 2.75±0.12 hours. Average recording duration was 2.85±0.06 hours (awake rest) and 17.67±0.44 hours (SWS). Brains were then sectioned to verify recording locations (Fig. 1F; Fig. S2A; Table S1) and virus expression, which did not differ across groups (t=0.276, p>0.05, unpaired t-test; Fig. 1D; Fig. S2B). Recordings were randomly interleaved between groups as the experimenters were blind to the group identity.

Spike sorting (*21*) yielded single units (CT: 176.86±11.82, 124.0±38.28; ET: 206.0±20.58, 118.72±36.85 for probe1 and probe2, respectively; Fig. 2A, Table S1). Putative excitatory (mean firing rate: 2.523±0.049 Hz) and fast-spiking (FS) interneurons (mean firing rate: 10.038±0.359 Hz) were classified based on the waveform and auto-correlogram features (Fig. S2D). Four animals were excluded due to low cell yield or surgical issues, resulting in a final sample of n=14 animals (CT: 7 (4M/3F); ET: 7 (4M/3F)) for the RSC probe and n=10 animals (CT: 5 (3M/2F); ET: 5 (3M/2F)) for the V1 probe.

**Fig. 2.**
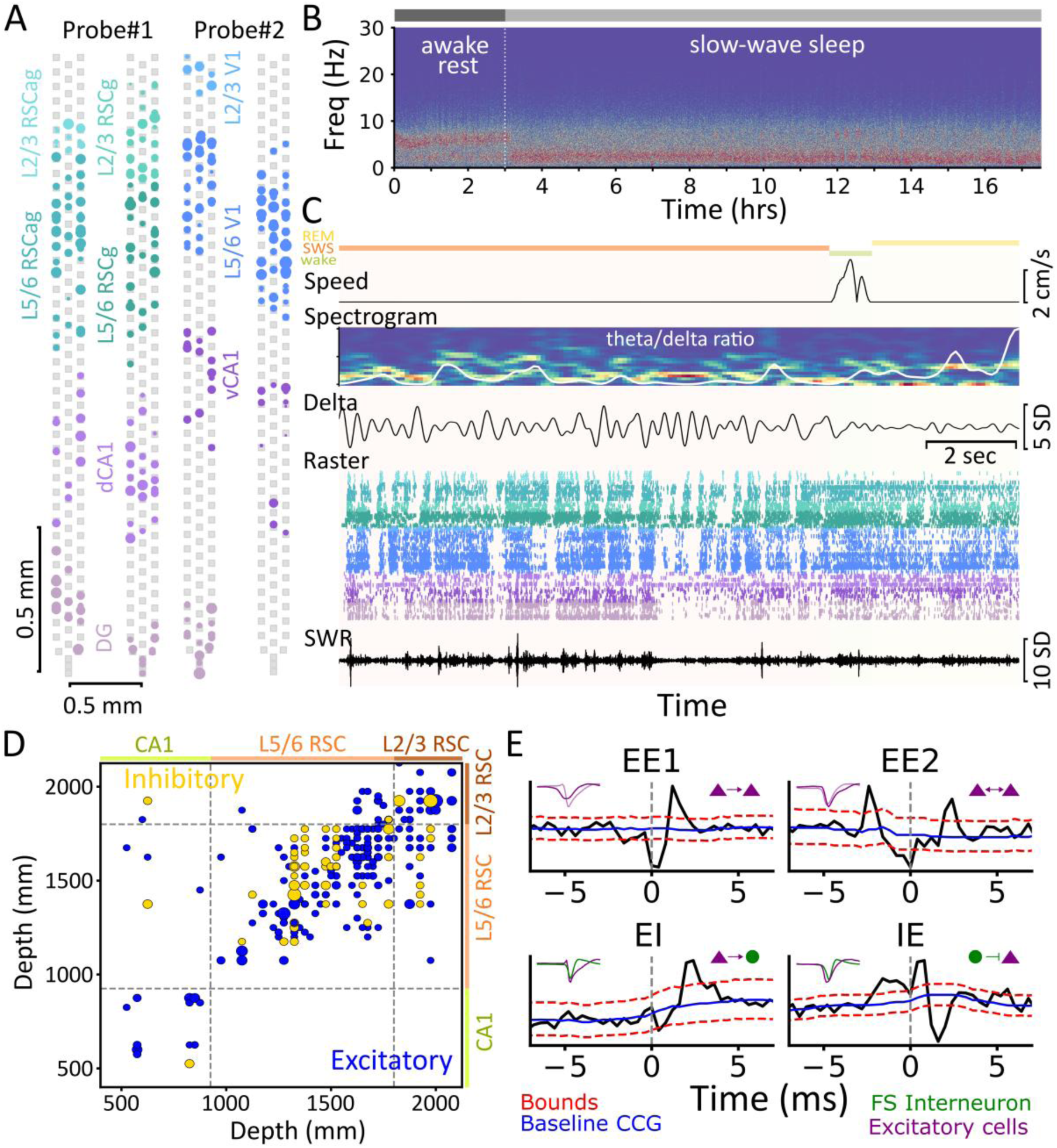
High-density electrophysiology recordings and monosynaptic connectivity analysis. (**A**) Left: 256-channel probe (gray squares mark recording sites spanning 2.125 mm) targeting DG (lily), dCA1 (lavender), L5/6 RSCg (sea green), L2/3 RSCg (pale teal), L5/6 RSCag (fountain blue), and L2/3 RSCag (aquamarine blue) from an example animal. Right: second probe targeting L2/3 V1 (light sky blue), L5/6 V1 (dark sky blue) and vCA1 (lilac) (L4 units excluded). Circles show single units; size reflects mean firing rate. (**B**) Spectrogram from an L5 RSCag channel showing 3 hours of awake rest, followed by ∼17.5 hours of induced SWS. (**C**) Brain states: SWS (orange), REM (yellow), and wake (green) are on top. Zoom-in on a 15 s segment showing speed (cm/s), spectrogram (0-20 Hz) with theta/delta ratio (white line) overlaid, delta-filtered (1-4 Hz) signal from the same channel as (B), spike raster sorted by region and depth, and sharp-wave ripple (SWR, 110-250 Hz, middle) filtered trace from dCA1. Same color scheme as (A) is used for spike raster. (**D**) Putative excitatory (blue, include EE and EI) and inhibitory (gold) connections across regions and layers (dCA1, L5/6 RSCg/RSCag, and L2/3 RSCg/RSCag) for a representative animal. Dot size indicates synaptic strength. In dCA1, excitatory connections observed are excitatory-to-inhibitory (EI) connections only. (**E**) Example Cross-correlogram (CCG) example showing unidirectional excitatory-excitatory (EE1), bidirectional excitatory-excitatory (EE2), excitatory-inhibitory (EI), and inhibitory-excitatory (IE). Excitatory units (E) and fast-spiking interneurons (I) are shown as purple triangles and green circles, respectively. Inset: waveforms of pre- and post-synaptic units. Solid blue line: baseline; red dashed line: confidence intervals (α=0.001).

Brain state (wake, SWS/NREM, REM) was scored using running speed, theta (5-10 Hz), and delta (1-4 Hz) oscillation power (Fig. 2B-C) (*22*). ET group showed slightly reduced delta power in the last 2-3 hours (F=1.741, p=0.023, group*time effect, rmANOVA) (Fig. S3B). Therefore, we analyzed only the first 15 hours (∼12 hours SWS). No group differences were found in total awake: CT: 2.56±0.12, ET: 2.6±0.11 hours (t=−0.185, p=0.85, unpaired t-test), SWS duration: CT: 9.9±0.22, ET: 10.1±0.2 hours (t=0.688, p=0.504; Table S1), SWS proportion (F=0.007, p=0.935; Fig. S3C), SWR rate (F=0.045, p=0.835, group-effect rmANOVA; Fig. S3C1), and other SWR-related properties (p>0.05, unpaired t-test with Bonferroni correction; Fig. S3C2-4).

### Changes in functional monosynaptic connectivity

We first analyzed functional synaptic connectivity in NC to test whether enriched experiences remodel NC circuitry to support encoding of large number of experiences. We computed cross-correlogram (CCG) for all simultaneously recorded neuron pairs and identified short-latency peaks or troughs (0.8-4.8 ms) to identify putative monosynaptic excitatory or inhibitory connections (*19*, *23*) (Fig. 2D). While we refer to these connections as functional, their validity is supported by juxtacellular recordings, which have demonstrated that such CCG peaks and troughs reflect true monosynaptic connections (*24*). Each significant connection was further classified as unidirectional excitatory-excitatory (EE1), bidirectional excitatory-excitatory (EE2), excitatory-inhibitory (EI), and inhibitory-excitatory (IE), based on short-latency peak/trough and cell classification (Fig. 2E). Common input connections with zero-lag synchrony, typically observed between interneuron pairs (II), were excluded (*24*).

We analyzed a total of 7,611.43±955.98 excitatory (EE1+EE2) and 3,818.43±525.23 inhibitory (EI+IE) cell pairs in RSC (n=7CT/7ET), and 3,376.5±710.25 excitatory and 1,880.5±344.81 inhibitory pairs in V1 (n=5CT/5ET). The total number of excitatory-to-excitatory (EE1+EE2) connections were comparable between groups in both RSC and V1. However, ET mice showed a significant increase in EE2 connections (t=−3.563, p=0.009; unpaired t-test; Table S2) and IE connections (t=−2.933, p=0.0158; Table S2) specifically in RSC, but not in V1 (Fig. 3A,B). This increase was accompanied by a significant decrease in EE1 connections in RSC (t=3.361, p=0.006; Table S2). No difference was observed in connection probability for EI connections.

**Fig. 3:**
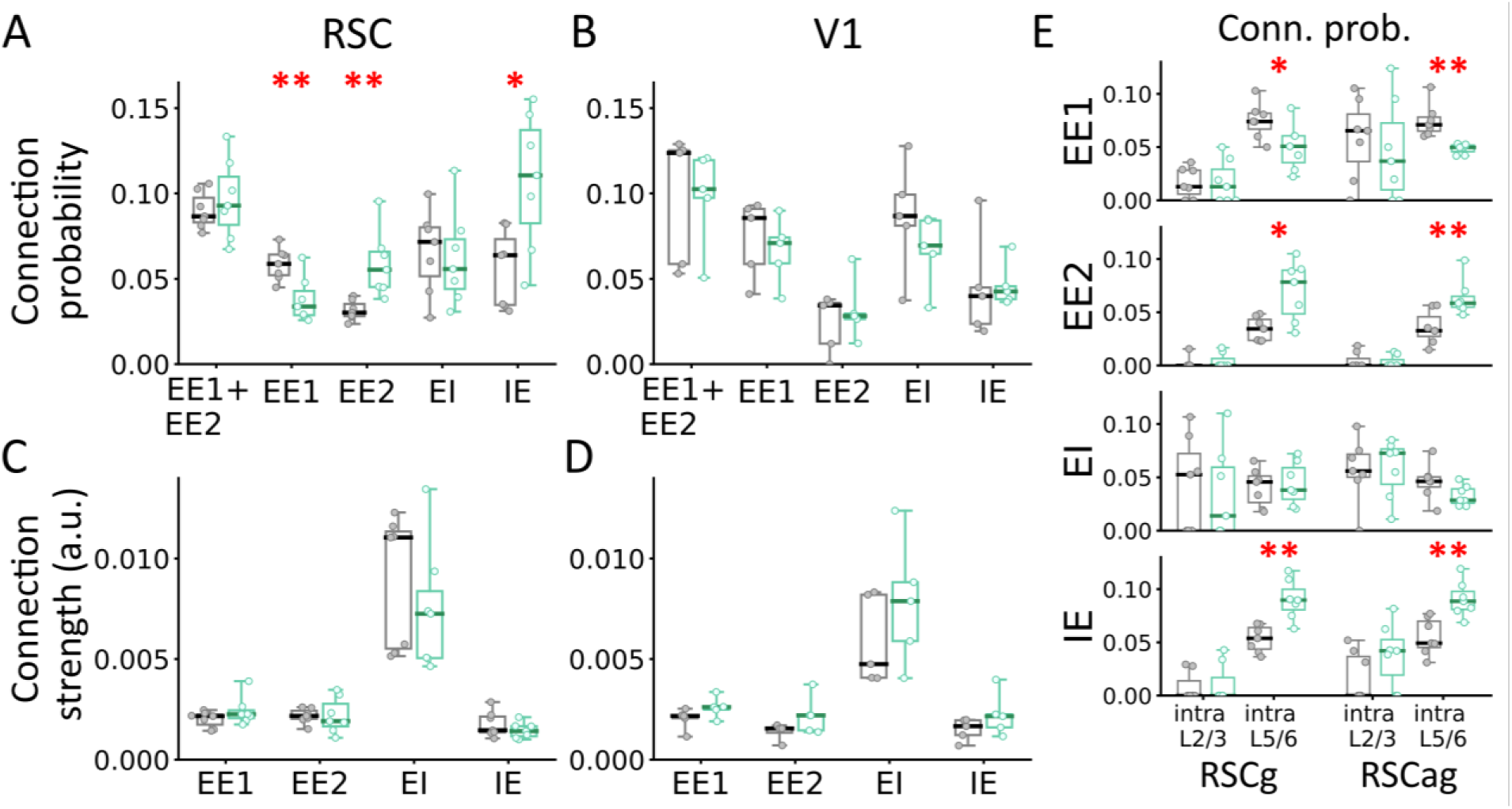
Enriched experiences alter functional monosynaptic connectivity in deep RSC. Connection probability for all excitatory-to-excitatory (EE1+EE2), EE1, EE2, and IE connections for (**A**) RSC (n=7CT/ 7 ET) and (**B**) V1 (n=5CT/ 5ET) for ET (aquamarine) and CT (gray) mice. Horizontal solid lines indicate the distribution mean. A similar plot is shown for synaptic interaction strength (a.u.) calculated from sum of values above or below chance CCG in the 0.8-4.8 ms range for (**C**) RSC and (**D**) V1 for both groups. (**E**) Connection probability for within-layers (intralaminar) EE1, EE2, EI, and IE connections (top to bottom) for RSCg and RSCag. P-values for unpaired t-test are listed at the top of each combination. *p<0.05, **p<0.01.

Next, we assessed synaptic connection strength using: 1) sum of observed CCG values exceeding (or below) chance, normalized by the reference cell’s spike count, 2) z-score peak (or trough) relative to the chance-level CCG. For EE2 connections, strength was averaged across both sides of the CCG. No significant group differences in connection strength were observed for any connection type in either region (Fig. 3C,D; Table S2; Fig. S4A). Consistent with previous studies, connection probability declined with distance (Fig. S4B). EI connections were generally stronger than EE1 and EE2 connections (Fig. S4B), and all connection types showed lognormal strength distributions (Fig. S4C) (*23*, *25*, *26*). Unlike a previous study (*23*), we did not observe stronger EE2 connections relative to EE1 (Table S2). No sex differences were observed (p>0.05, ANOVA with Bonferroni correction, Fig. S7A).

The expected number of unconnected, EE1, and EE2 pairs was estimated using the following equations: N(1-p)^2^, 2Np(1-p), and Np^2^, respectively, where N is the total EE pairs and p is the overall EE connection probability (*19*, *23*). Using p=0.1 (p_EE1_ + p_EE2_; Fig. 3A,B) and the observed N, we found that the number of EE1 connections were lower than expected, whereas EE2 were significantly higher (∼3x) than expected (p<0.05). Similar increase in EE2 connections have been reported previously (*19*, *27*). This overrepresentation of bidirectional EE2 (symmetric) connections is consistent with attractor dynamics and may reflect circuit-level changes supporting the encoding of multiple items experienced during ENR (*5*).

In V1, we analyzed intralaminar deep (L5/6) connections in Fig. 3B,D, as insufficient number of superficial (L2/3) neurons were available across animals. In contrast, RSC data included both L2/3 and L5/6 neurons from granular (RSCg) and agranular (RSCag) subdivisions. Therefore, we decided to analyze connections within and between L2/3 and L5/6of RSCg and RSCag. Interareal connections were rare, but we detected some interlaminar connections within the same brain region for EE1 (probability in RSCg: CT: 0.011±0.003, ET: 0.012±0.004; RSCag: CT: 0.025±0.004, ET: 0.022±0.005), EI (RSCg: CT: 0.015±0.004, ET: 0.016±0.006; RSCag: CT: 0.020±0.005, ET: 0.020±0.007), and IE (RSCg: CT: 0.019±0.004, ET: 0.016±0.004; RSCag: CT: 0.008±0.004, ET: 0.010±0.005) connection types, but not for EE2, with no significant group differences in their probability or strength. Thus, we focused on intralaminar connections within each RSC subregion.

The total number of intralaminar EE pairs were: L2/3 RSCg: 91.08±17.89; L5/6 RSCg: 1194.0±211.01; L2/3 RSCag: 82.64±10.56; L5/6 RSCag: 1417.21±176.36 and EI/IE pairs were: L2/3 RSCg: 71.63±11.5; L5/6 RSCg: 620.93±118.3; L2/3 RSCag: 70.69±13.44; L5/6 RSCag: 454.21±70.48. In ET mice, EE2 connection probability was significantly higher in deep (L5/6) layers of both RSCg (t=−3.149, p=0.013) and RSCag (t=−3.177, p=0.008) compared to CT mice (Fig. 3E). In contrast, the EE1 connection probability was significantly lower in ET mice in L5/6 of both RSCg (t=2.36, p=0.037) and RSCag (t=4.259, p=0.003), consistent with Fig. 3A results. EI connection rates were unchanged across groups, but IE connection probability was significantly increased in ET mice in L5/6 of both RSCg (t=−4.337, p=0.0013) and RSCag (t=−3.894, p=0.002) (Fig. 3E). Across animals, L2/3 connection probabilities were often zero, suggesting that the results in Fig. 3A are largely driven by L5/6 neurons. Finally, no group differences were observed in intralaminar synaptic strength across connection types in either RSCg or RSCag (Fig. S4D). Subsampling L5/6 neurons to match the number of L2/3 neurons yielded similar results, ruling out sampling bias.

We computed CCG over the entire 15-hour recording to obtain a reliable estimate of connection probability (*19*). However, awake rest and SWS differ in several aspects, including activity levels, neuromodulatory tone, memory replay dynamics, etc. (*28–31*). To assess state-dependent changes, we recalculated connection strength for connections identified in the full recording, separately for awake rest (∼2.5-3 hours) and the first 3 hours of SWS. Previous studies reported that intralaminar L5/6 EE connection strength increases from awake rest to SWS (*25*). We also observed the largest differences in intralaminar L5/6 EE and IE connection probabilities (Fig. 3A,B). We therefore focused our analysis on intralaminar L5/6 connections to determine whether similar changes in connection strength occur. No significant group differences were found; although EI, IE, and EE2 connections showed a slight upward trend from awake rest to SWS in both groups (Fig. S4E), while EE1 remained stable. Together, these results highlight selective reorganization of deep-layer RSC connectivity following EE, characterized by a shift toward more reciprocal (bidirectional) excitatory connections and enhanced inhibitory feedback, consistent with attractor-like dynamics.

### Changes in population coding dynamics

Having established the changes in synaptic connectivity, we next asked whether ET mice would show sparser and more orthogonal population activity. We computed population and lifetime sparseness using 50 ms binned population vectors (PV) of excitatory units during awake rest and SWS. Lifetime sparseness reflects how selectively a single neuron fires over time, while population sparseness quantifies the proportion of active neurons in a population at any given time (*32*, *33*). Neuron counts were balanced across animals and brain regions by sub-sampling 1000 times and averaging. We compared units from CA1, L2/3 RSC, and L5/6 RSC separately due to their distinct coding properties (*34–36*). For V1, we focused on L5/6 neurons due to insufficient L2/3 neurons.

During both awake rest and SWS, ET mice showed significantly increased population and lifetime sparseness in L5/6 RSCg (lifetime sparsity: Awk Rest: t=−3.147, p=0.012; SWS: t=−6.628, p<0.001; population sparsity: Awk Rest: t=−3.23, p=0.008; SWS: t=−4.897, p<0.001) and L5/6 RSCag (lifetime sparsity: Awk Rest: t=−3.812, p=0.004; SWS: t=−4.8, p<0.001; population sparsity: Awk Rest: t=−2.36, p=0.038; SWS: t=−3.15, p=0.009) (Fig. 4A,B; Table S3). A modest but significant increase in population sparseness was also observed in L2/3 RSCag in ET mice during SWS (t=−2.445, p=0.043; unpaired t-test). No group differences in sparseness were found in other regions (Fig. 4B, Table S3). Dividing the 12-hour SWS into 3-hour blocks showed comparable results, with ET mice exhibiting increased sparsity in RSC L5/6 neurons (Fig. S5A,B). As no systematic differences were observed across SWS blocks, we report data pooled across the entire SWS period (Fig. 4).

**Fig. 4:**
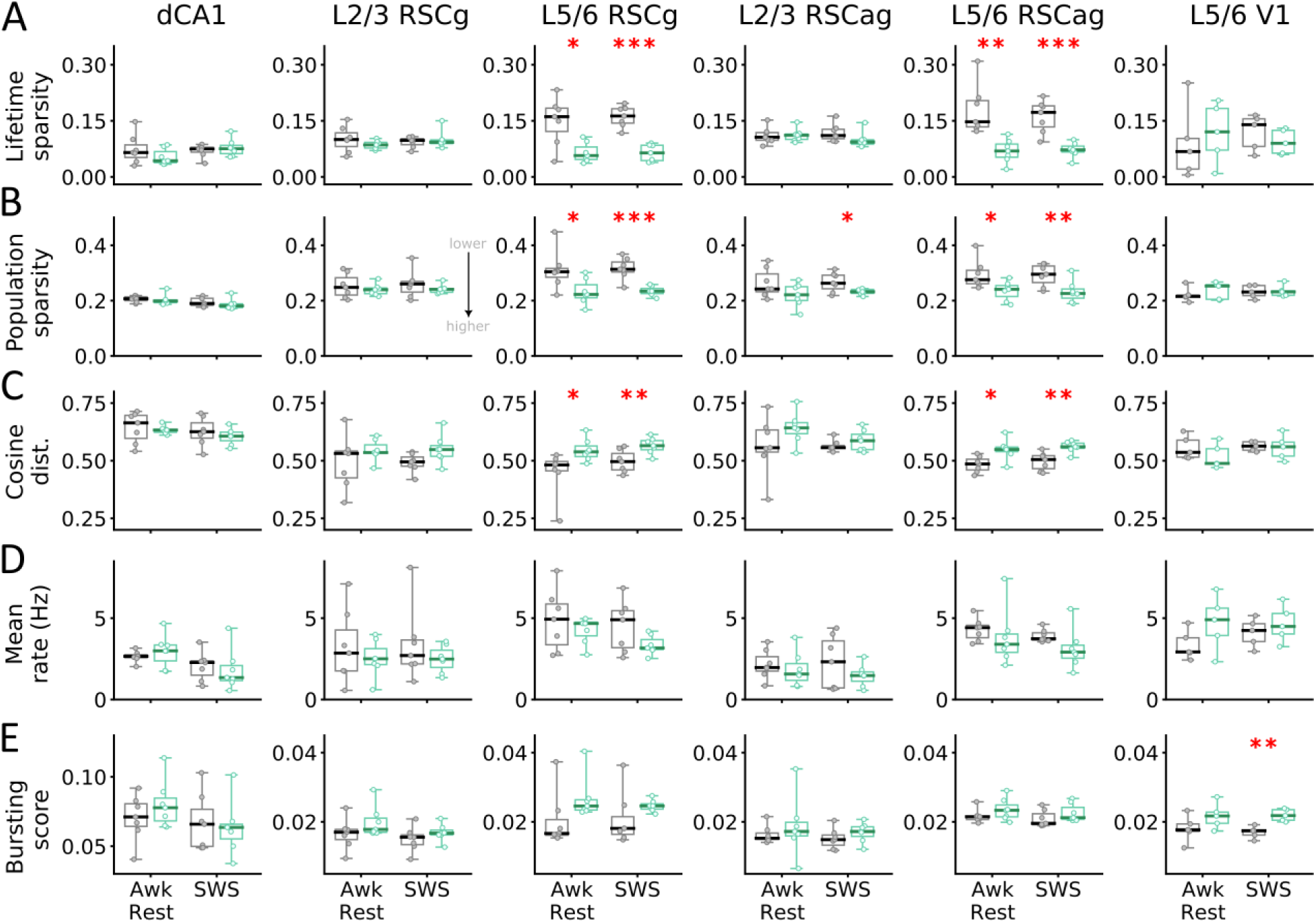
Enriched experiences increase sparsity and orthogonality in RSC deep layers. Changes in (**A**) lifetime sparsity, (**B**) population sparsity, (**C**) cosine distance, (**D**) mean firing rate (Hz), and (**E**) bursting score for excitatory neurons across brain regions: dCA1, superficial (L2/3) and deep (L5/6) granular and agranular retrosplenial cortex (RSCg/RSCag) and primary visual cortex (V1) for ET (aquamarine) and CT (gray) mice during awake rest and SWS. Lower values indicate more sparsity (A,B). Each point represents data from one animal. Total n=14 mice (7CT/ 7ET) for RSC and dCA1, and n=10 (5CT/5ET) for V1. ET mice show higher lifetime, population sparsity, and cosine distance for L5/6 RSC ensembles, but not for V1. *p<0.05, **p<0.01, ***p<0.001.

We then looked at orthogonality across brain regions using average cosine distance between pairs of PVs. ET mice exhibited significantly larger distances in L5/6 RSCg (Awk Rest: t=−2.816, p=0.017; SWS: t=−3.15, p=0.009) and L5/6 RSCag neurons (Awk Rest: t=−2.725, p=0.023; SWS: t=−3.464, p=0.005) (Fig. 4C, S5C; Table S3), indicating a more diverse neural state space. Other regions did not show significant group differences. Increase in sparsity and orthogonality were generally more pronounced during SWS, potentially reflecting differences in activity levels and memory replay dynamics between the two states (*28–31*). Perhaps a more diverse activity space is being explored during SWS than awake rest, which might be more prominent in ET mice. Differences in sparsity and orthogonality could not be fully explained by differences in mean firing rate or bursting index (proportion of spikes with inter-spike interval < 6 ms), as no significant group effects were detected for either measure (Fig. 4E,F, Fig. S5, Table S3). That said, a trend toward more bursting and lower firing rates in ET mice was visible. Consistent with previous studies, mean firing rates were generally higher in NC L5/6 compared to L2/3 (*34–36*) and lower during SWS than awake rest (*37*, *38*). No significant sex differences were found (p>0.05, ANOVA with Bonferroni correction, Fig. S7b).

Changes in population and lifetime sparsity were primarily observed in L5/6 RSC. The lack of significant changes in CA1 and L2/3 RSC may be attributed to these regions already operating at near-maximal (or near-optimal) sparsity levels, whereas L5/6 neurons may have greater potential for improving coding efficiency (*34*, *35*). No significant difference in lifetime sparsity was observed in DG neurons after pooling data across animals (Fig. S6A). This finding, combined with the lack of difference in CA1 sparsity, contrasts with previous studies examining hippocampal subregions after ENR (*39–41*). As CT mice ran more laps (Fig. 1C), more physical exercise may have promoted neurogenesis and hippocampal sparsity, potentially reducing differences between the groups (*42*, *43*). L2/3 RSC likely follows a similar pattern, as it receives direct inputs from dCA1 (*44*, *45*). However, a more likely explanation is that L5/6 neurons have been shown to code for categorical knowledge (*46*, *47*). Consequently, larger effects are expected in L5/6 of ET mice to accommodate a greater diversity of sensory-motor experiences. Selective increase in L5/6 RSC, but not in L5/6 V1, may result from V1 being an early sensory area where feature coding is relatively less experience-dependent. Alternatively, RSC integrates multimodal inputs from multiple areas beyond V1, such as somatosensory and motor, which may exhibit more pronounced changes and drive the observed results (*48*). Supporting this, a recent study using the same ENR paradigm demonstrated rapid stabilization of spatially selective firing in secondary motor cortex, a primary input region to RSC (*11*). Further exploration of PV correlations using graph-based analysis trends towards lower clustering coefficient (not significant) for L5/6 units in ET mice, indicative of weaker coactivation, in alignment with higher orthogonality observed (Fig. S6C).

## Discussion

The most significant finding of this study is the *in vivo* validation of a key prediction from auto-associative attractor networks: that knowledge accumulation should lead to an increase in reciprocal (bidirectional) excitatory-excitatory connections (*1–4*). Our results demonstrate that enriched experience, used here as a model for knowledge accumulation, produces a substantial shift from unidirectional to bidirectional excitatory-excitatory connections in higher-order cortex, providing key experimental evidence for this long-theorized mechanism of cortical rewiring. This finding has important implications for our understanding of how the brain stores and organizes knowledge. Attractor network models predict that reciprocal synaptic connectivity enhances robustness, enables signal amplification, and supports pattern completion (*5*), all critical functions for maintaining stable memory representations. The observed shift toward more symmetric excitatory connections in ET mice, exposed to a wide range of experiences, directly supports these theoretical predictions and suggests that enriched brains develop connectivity patterns optimized for storing multiple, stable attractor states.

Complementing this primary finding, we observed that ENR also enhanced representational sparsity and orthogonality in population codes, particularly in deep RSC layers. ET mice exhibited significantly increased population and lifetime sparseness in L5/6 RSC during both awake rest and slow-wave sleep, accompanied by larger cosine distances between population activity. We also found a trend toward increased bursting and a reduced mean firing rate in L5/6 RSC (though not significant), suggesting that their combined effect may contribute to the observed changes in coding sparsity. These changes in sparsity align with network simulations, which show that increased sparsity and bidirectionally connected excitatory pairs characterize systems optimized to maximize storage capacity through fixed attractors (*23*, *49–51*). Together, the convergence of our connectivity and population coding results provides compelling evidence that ENR reorganizes cortical circuits according to principles predicted by computational theories of memory storage.

The observed increase in inhibitory-to-excitatory connections could serve to balance the high gain resulting from more bidirectional excitatory connections, thereby preventing runaway excitation. Alternatively, consistent with modeling work showing that enhanced inhibition promotes the formation of non-overlapping cell assemblies (*52*), increased inhibition in ET mice may facilitate the development of orthogonal cell assemblies, reflected in the prevalence of bidirectional excitatory connections and greater sparsity. The sparser and more orthogonal representations we documented may promote faster learning by minimizing overlap between representations, thereby reducing interference with existing memories (*53–55*). This could contribute to the enhanced behavioral and forward knowledge transfer consistently observed in enriched animals (*8*, *10*, *11*).

Several limitations should be acknowledged. We used a specific ENR paradigm (*10*) that significantly improves behavioral performance compared to standard protocols, but our findings may not generalize across all enrichment paradigms. The chemogenetic enhancement of SWS (*18*, *56*), while enabling long, stable recordings necessary for detecting weak excitatory connections (*19*), disrupted normal sleep cycles during recording sessions (which were conducted long after the enrichment treatment itself). Future work should examine how connectivity changes evolve over the course of enrichment protocol. In addition, assessing whether comparable changes occur in other cortical regions, such as prefrontal, motor, or somatosensory cortex would help establish the generality of our findings. It would also be interesting to compare higher-order visual and somatosensory cortex in modality-specific enrichment, i.e., visual-only or somatosensory-only.

In summary, our findings provide evidence supporting cell assembly (*1*) and modern attractor network predictions, including Hopfield networks, (*2–4*) regarding cortical connectivity changes following knowledge accumulation. The observed shift toward enhanced bidirectional excitatory and inhibitory connections, accompanied by sparser and more orthogonal population representations, suggests that enriched experiences reorganize cortical circuits to optimize memory storage and retrieval. These results link theoretical predictions from computational neuroscience with empirical observations, advancing our understanding of how lifelong learning shapes brain connectivity and function.

## Acknowledgments

The authors thank Scott Kilianski for helping with virus injections, Priscilla Ee for helping with surgeries and histology, and Meenakshi Chandrasekaran, Varleen Kaur, James Musasizi, Yinzhi Ying, Tianyi Tong, Yimin Lei, and Kaelyn Chen for helping with running enrichment behavior.

## Author contributions

Conceptualization: BLM, RS

Methodology: RS, JLS, CA, BLM

Formal analysis: RS

Investigation: RS, JLS, WN, AAA

Data curation: RS

Writing-original draft: RS, JLS, BLM

Writing-review & editing: RS, BLM, JLS, AAA, WN, CA

Visualization: RS

Supervision: BLM

Project administration: BLM

Funding acquisition: BLM

## Data and materials availability

All processed data and analysis scripts will be made public before publication.

## List of Supplementary Materials

Materials and Methods

Figs. S1 to S7

Tables S1 to S3

## Materials and Methods

### Animals

All experiments were approved and conducted in accordance with the guidelines set by the Institutional Animal Care and Use Committee (IACUC) of the University of California, Irvine. Mice were housed in pairs on a 12-hour on/off light cycle and were not food-deprived throughout the experiment. N=18 (10M/ 8F) Vgat-Cre transgenic mice (*Slc32a1^tm2(cre)Lowl^*, Jackson Laboratory) were used for the experiment. These animals express Cre recombinase in Vgat-expressing cells, primarily GABAergic inhibitory neurons. Four animals were eventually excluded from analysis due to low cell yield, poor virus expression, incorrect recording location, or death during surgery, resulting in a final sample of n=14 animals (CT: 7 (4M/3F); ET: 7 (4M/3F)) for RSC and n=10 animals (CT: 5 (3M/2F); ET: 5 (3M/2F)) for V1.

### Environmental Enrichment (EE) Protocol

We used an enrichment protocol similar to that described in Gattas et al. (*10*), with certain modifications listed below. Enrichment (ET) and control (CT) tracks (track dimension: 86.5cm long x 86.5cm wide; arm width: 11.43cm) were placed next to each other, separated by a large opaque wall to block the view of the other track. Each arm on the track also had large transparent walls, both inside and outside of it, except the common opaque wall (Fig. 1B). A top-down camera recorded laps completed, lap duration, and reward delivery. To prevent backtracking, one-way doors were installed at the first and third corners of the track. The training protocol started at the age of two months. Mice were first handled for 15-min for one week. Handling was paired with sweetened milk reward. Throughout training, animals were maintained on an *ad libitum* diet and were self-motivated to run, sometimes not consuming the reward. Animals ran at fixed times daily, and their weight was noted at the start of each session. No significant difference in body weight was noticed between both groups (p>0.05). In an individual session, two cage-mates (1ET/ 1CT) ran simultaneously on their respective tracks for 1-hour per session, five days per week, for 10–12 weeks. This differs from Gattas et al. (*10*), where ENR was run six days per week for 8-9 weeks. ET mice ran in clockwise direction, while CT mice ran in counter-clockwise direction. Mice received a sweetened milk reward upon completing each lap.

During the first week of habituation, both groups were exposed to the track with only ramps (3 per arm, 12 in total, 20 cm long x 11 cm wide) and one-way doors. CT had the same ramps for the entire enrichment/control period. On the other hand, the ET had three obstacles per arm and a total of twelve unique obstacles (except for the habituation period, where all arms had ramps) (Fig. 1B, Fig. S1A). After the first week, ET mice were introduced to new obstacles (replacing one ramp in each arm per day) until distinct obstacles replaced all ramps. Once obstacles replaced all ramps, we made changes to ensure new obstacle configurations daily. This was done by either replacing old obstacles with new ones or changing the arrangement (order) of old obstacles by moving them to new locations or rotating them (Fig. S1A,C). Three changes were made in each 1-hour session at 20-min. intervals. The daily changes to the obstacle course provided the ET group with a diverse array of sensory-motor experiences.

The CT group acted as a control for exercise, as exercise can influence population activity and behavior performance (*57*). CT animals ran more laps than the ET (Fig. 1D) (F=5.938, p=0.033, group-effect, rmANOVA). We also recorded the time taken to finish each lap in each session. As expected, ET animals typically took longer to finish each lap.

### Behavior tests

Behavioral performance was assessed using a hippocampus-dependent spatial y-maze test (*58*). A separate cohort of n=15 (8CT/7ET) mice were exposed to a y-maze with one arm blocked (counter-balanced) for 10 min. (arm length: 8.9 cm wide and 26.67 cm long) The y-maze had transparent walls, so nearby objects (< 15 cm distance, 15-25 cm height) were visible from other arms and multiple distal room cues. After 24-hours, mice were allowed to explore the y-maze with the closed arm open for 5 min (Fig. S1C). Animals were placed at the beginning of the start arm facing forward for training and test sessions. We calculated the Discrimination ratio 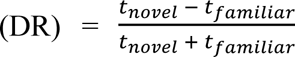, measuring only active moving time. Surprisingly, both groups showed no difference (p=0.285, unpaired t-test) and were comparable to baseline (p>0.05) (Fig. S1C,F).

We hypothesized that the transparent walls and objects nearby may have enabled both groups to form a unified context representation, affecting the results. To test this, we conducted an additional experiment where the novel arm and its associated object were visually blocked (blocked y-maze) during training and testing. This task was performed with the same parameters as the spatial y-maze above but included a rotated and translated maze, different floor textures, and new objects nearby. Importantly, barriers were added to hide the novel arm and its associated objects (Fig. S1D,F). ET mice showed better discrimination than the CT mice (t=−4.2, p=0.0013, unpaired t-test) (Fig. S1F). The improved performance in ET mice after blocking the novel arm supports our previously stated hypothesis, thereby highlighting the superior spatial recognition memory of ET mice, consistent with previous findings (*10*). Note that in the original ENR paper (*10*), experiments was done similar to the blocked y-maze.

We then assessed view-based recognition using the view-invariant object recognition (VIOR) task. This test examines whether animals recognize rotated vs. non-rotated objects as familiar and explore them equally. During training, animals explored a y-maze with opaque walls (no other nearby object, unlike previous tasks) for 10 min, encountering the same object (a green frog toy) in the same orientation in two arms other than the start arm (Fig. S1E,F). After a 5 min break, mice were reintroduced to the maze for 3 min, where one of the objects was rotated by 90^◦^ (counter-balanced arm). The objects and behavior apparatus were wiped with 70% ethanol to eliminate lingering odor cues. Both groups spent equal time exploring both arms (unpaired t-test, p=0.093), suggesting that they treat both objects as equally familiar (Fig. S1F). Our results show a similar trend as previous study (*10*), i.e., better performance in the ET group, but no statistial significance. Repeating the test with increased difficulty (longer gap between test and training session) and a larger number of animals should improve the results.

### Surgical Procedures

*Headbar + Virus injection surgery:* After the enrichment/control protocol (∼5.5 months old), animals were handled for five days, then implanted with headbars and injected with *pAAV2-hSyn-DIO-hM3D(Gq)-mCherry* virus to induce slow-wave sleep (SWS) (*18*, *56*) (Fig. 1D). The virus expresses Cre-dependent excitatory *hM3DGq* DREADD receptors under the hSyn promoter, and mCherry is a fluorescent protein. After leveling the skull and marking bregma and lambda, we used a pump-driven Hamilton syringe (Nanoliter2020, WPI, FL, USA) filled it with 3 µL distilled water, 0.2 µL air bubble, and 3 µL virus to inject 100-125nL of virus bilaterally into the Parafacial Zone (PZ) (AP-5.3 mm, ML±0.7 mm, and DV-4.3 mm from bregma; DV is from the top of the skull). The injection was done at 30-40 nL/min, and the syringe was left in place for 5 min before removal. The needle was checked for blockage before and after insertion. Injection site burr holes were sealed with bone wax and a layer of Metabond (Parkell, NY, USA). Stainless steel headbars were then cemented to the skull, with ground and reference pins implanted over the cerebellum (Fig 1E). Post-surgery, animals were allowed a 7-day recovery period and provided with ibuprofen (2 mg/mL) and amoxicillin (0.5 mg/mL) in their drinking water. After recovery, we started head-fixation habituation and did one-shot electrophysiology recording (see below).

*Craniotomy and catheter implant surgery:* Approximately 2.5 months after headbar surgery, once animals were acclimated to resting on an air-suspended styrofoam ball, a craniotomy was performed over target sites in the left hemisphere (except for n=3 animals in right hemisphere) determined using the Allen mouse brain atlas (*59*, *60*). Craniotomy was performed to simultaneously record from dorsal CA1 (dCA1), granular Retrosplenial cortex (RSCg), agranular Retrosplenial cortex (RSCag) (medial shank: AP-1.8 mm, ML-0.8 mm) and primary visual cortex (V1), and ventral CA1 (vCA1) (medial shank: AP-3.2 mm, ML-2.8 mm) in the left hemisphere (Fig. 1F). In 4 animals, recordings were done from site #1 only, whereas dual-site recordings were performed in the remaining animals. In 3 animals (2ET/1CT), craniotomy was performed over the right hemisphere. After craniotomy, the exposed skull was sealed with sterile Vaseline, Kwik-Cast sealant (WPI, FL, USA), and Metabond. Note that the dura was left intact. Finally, a catheter was implanted to deliver Clozapine N-oxide (CNO). The catheter was made of 0.4 mm diameter tubing (suitable for fitting a 28 G needle) and placed under the mice’s upper back skin, with only a 1-mm section inside. The other end of the catheter was cemented to the opposite hemisphere headbar to prevent interference with probe insertion (Fig. 1F). The catheter was checked for leakage by running saline through it. Post-surgery, animals were monitored until active, then returned to their home cages.

### Experimental Design

Two-month-old mice were first handled for a week. These pair-housed mice were then run on an automated side-by-side enrichment/control setup, such that one mouse ran on CT while its cagemate ran on ET for 10-12 weeks, with 5 x 1-hour sessions per week. Mice received a sweetened milk reward at the end of each lap and were not food-deprived. ET mice had to navigate through twelve different obstacles (new obstacle configuration every session). In contrast, CT mice ran on simple ramp hurdles that remained the same throughout the ENR protocol (Fig. 1B). We recorded the weight, number of laps, and time taken to finish each lap. For the remainder of the experiment, the personnel involved were blind to the group identity.

Following the ENR protocol, mice were trained to be head-fixed for acute electrophysiology recordings. To examine population activity state space and synaptic connectivity during offline periods, such as awake rest and slow-wave sleep (SWS), mice were implanted with headbars and injected with pAAV2-hSyn-DIO-hM3D(Gq)-mCherry virus bilaterally into the Parafacial Zone (PZ) to induce SWS (*18*) (Fig. 1D,E). This approach allowed us to record long-duration resting data, which is typically hard to achieve in head-fixed animals on a running ball. The longer recording duration also improved our capability to detect long-range excitatory-excitatory (EE) connections across layers and brain regions (*19*). Following a week of recovery, mice were gradually trained to be head-fixed on a suspended running ball (80-cm circumference), starting from 15-min and increasing to 2-hours over ∼2 months. The head-fixed training was done in a dark, silent room. Three tablets with black screens covered the animal’s field of view. A sweetened milk reward was delivered every 10 min via a lick tube. We tested the success of virus injection at the 1-month and 2-month mark from surgery for each animal by injecting CNO (subcutaneously 0.5 mg/kg) and confirming if the animal fell asleep (lost mobility). For this, we took a paired ET/CT group and randomly injected one animal with CNO and another with saline. A side camera tracked their movement inside the cage (lid left open) to confirm sleep induction. After virus injection and head-fixed training, we performed craniotomies over RSC, V1, DG, and CA1 (dorsal and ventral) and implanted a catheter in the upper back to inject CNO subcutaneously (Fig. 1E,F).

The day after craniotomy surgery, we recorded single-unit and local-field potentials (LFP) data for 2.5-3 hours of awake rest, followed by at least 13-hours of induced SWS data for each animal using dual 256-channel silicon probes (*20*). Multiple booster shots of CNO (0.5 mg/kg) were given every 2.75±0.12 hour via catheter to keep the mice in continuous SWS. Recording was done in the same room as training, with lights off and tablets with black screens. Recordings started between 5:30-7:00 PM across animals. We maintained the room temperature at 37°C using a room heater. The total recording duration across animals for awake rest was: 2.85±0.06 hours and induced SWS: 17.67±0.44 hours. Recordings were interleaved between ET and CT. Mice were perfused, and their brains were extracted at the end of the recordings to track probe marks and check virus expression.

### Silicon probe recordings

High-density silicon probes were used for the acute recordings (*20*). We used dual 256-channel probes (total 512 channels) targeting the RSCg, RSCag, dCA1 (probe#1), and V1, vCA1 (probe#2). Only probe#1 was used for n=4 animals (2ET/ 2CT). Each 256-channel probe (2 shanks separated by 0.5 mm, 128 channels per shank) had a vertical span of 2.125 mm with 25 µm vertical and 20 µm horizontal spacing between contacts arranged in a staggered grid-like pattern. The wideband electrophysiology data were acquired at 30 kHz and digitized at 16 bits on the headstage amplifier before being transmitted to the Intan acquisition system (Intan Technologies, CA, USA). Two separate digital sync pulses were also sent to the acquisition system and stored to mark 1) the times the animal moved on the running ball via rotary encoder or 2) a frame was acquired on the face tracking camera (recorded for only some animals and hence not analyzed). Intan RHX data acquisition software was used to acquire the data. The wideband data were low-pass filtered to extract LFP and resampled at 1000 Hz. Before each recording, the probe tips were coated with a fluorescent dye (Invitrogen DiD) and mounted on a 4-axis micromanipulator (Quad Sutter Instruments, Novato, CA, USA). Once the mouse’s head was leveled, the probe tip was used to measure bregma coordinates to decide the AP and ML coordinates for the target recording site. The probes were lowered at a speed of 1 µm/s until they reached the desired depth, with insertion monitored using a surgical microscope (Seiler Instruments, MO, USA). After reaching the target depth, sterile mineral oil was applied to the dura, and a settling period of 45-60 min was ensured before starting the behavioral task and neural data acquisition. Following the experiment, the probes were cleaned using distilled water, 70% ethanol, 1% Tergazyme, and a Trypsin (Invitrogen) protease solution.

### Histology

Following isoflurane, mice were transcardially perfused with cold 1X phosphate buffer solution (PBS), followed by 4% paraformaldehyde (PFA), and the extracted brain was stored in 4% PFA at 4°C. Before slicing, brains were transferred to 30% sucrose/PBS solution and stored at 4°C. We then coronally sectioned brains at 40 µm using a cryostat (Thermo Scientific HM525 NX) at −20°C, and each section was stored in well plates holding cryoprotectant solution. We selected brain slices with the deepest visible probe track marks (DiD staining) for immunohistology staining. We incubated slices while shaking at 4°C for 48 hours with primary antibodies: anti-NECAB1 (Atlas Antibodies, HPA023629) (*61*) at 200-fold dilution and anti-Wfs1 (Proteintech, 26995-1-AP) (*62*) at 8000-fold dilution to visualize layer 4 and layer 2b, respectively. Sections were washed with 1X tris buffered saline (TBS) and 0.1 M phosphate buffer and then incubated with secondary antibody Alexa Fluor 555 (Invitrogen, A-21428) at 400-fold dilution for 2 hours at room temperature. Slices were mounted onto slides with media containing DAPI (Invitrogen SlowFade Glass Soft-set Antifade Mountant with DAPI). A Keyence BZ-X1810 widefield fluorescence microscope with a 4x objective lens was used to acquire images, and the filter cubes DAPI, Cy5, and Texas Red were used to visualize nuclei, DiD silicon probe marks, and virus expression, respectively. The images were then aligned to the Allen Mouse Brain Atlas and Common Coordinate Framework (CCFv3) (*60*, *63*) and used to assign brain regions. Further analysis looking at the maximum power in the high frequency (> 300 Hz) (*25*), reversal of sharp-wave around CA1 cell layers (*64*), and large amplitude delta oscillation in the deep cortex (*25*) were used to confirm the brain region. Finally, we calculated the percentage of area with virus injections in PZ for AP-5.3, AP-5.7 mm, and AP-5.9 mm coronal sections and compared across animals. No significant difference was observed between groups (t=0.276, p=0.787, unpaired t-test) (Fig. 1D, Fig. S2B).

### Spike sorting and manual curation

Kilosort2.5 (*21*) was used to run spike-sorting. We used batch size=8 s from default 2 s for the drift estimation algorithm. Double-counted spikes were removed using the method described in (*65*). Briefly, we removed spikes within peak times within 5 data points (0.16 ms) and peak waveforms within five channels (around 50 µm) from Kilosort2.5 output. The maximum number of spikes extracted per batch and template-matching iterations were also increased compared to the default version. Manual inspection was performed using Phy to remove noise units and label units as MUA or good (based on ISI violations, waveform shape, etc.). Phy was also used to split over-merged clusters and merge over-split clusters.

### Unit metrics and quality control

We calculated various metrics based on a combination of the physical characteristics of the units’ waveforms and firing properties (*65*) for all good units labeled after manual inspection in Phy. Thresholds used for unit inclusion for final analysis: ISI violation rate < 0.2 (*66*), mean firing rate > 0.01 Hz (given long recordings), mean waveform amplitude > 30 µV, and presence ratio > 70%. No significant difference was seen between the quality metrics of units between CT and ET mice. Furthermore, we included measures of stability similar to the ones described in (*19*). Briefly, we calculated the fractional change in amplitude of each neuron in 3-hour blocks of induced SWS period (6-7 blocks depending on total duration). Only units with fractional change < 20% in both parameters were considered stable and included in the analysis. Similar analysis was done to ensure the overall population firing rate was stable during awake rest and SWS epochs. The epochs were considered separately due to lower mean firing rates in SWS (*22*, *38*).

### Cell-type classification

We first excluded non-negative units: positive and biphasic units (5.7±1.9%, Fig. S2D) corresponding to return currents and axonal spikes, respectively (*67*). The cell classification was performed using CellExplorer software using a combination of mean firing rate, waveform features, and auto-correlogram (acg) features (*68*) and further curated manually. Units with trough to peak duration of mean waveform<=0.425 ms were labeled as fast-spiking (FS) interneurons (mean firing rate: 10.038±0.359 Hz) and the rest as putative excitatory units (mean firing rate: 2.523±0.049 Hz). Comparable results were obtained after applying principal component analysis (PCA) on the features from waveform and acg of all units followed by k-means clustering.

### Spectral parameters calculation

We verified the placement of probes using LFP spectral analysis in addition to histology. We selected a series of vertical channels across the probe depth and calculated each channel’s relative power in the high-frequency range (100-500 Hz) (*25*, *69*). Across animals, we noticed a prominent peak in LFP power around cortex layer 5a and CA1 cell layer (Fig. S2C). Based on this, we selected a channel on the lateral shank targeting RSCag and V1 deep layer, which exhibited the maximum relative power, to extract relative power in different frequency bands: delta (1-4 Hz), theta (5-10 Hz), sigma (10-16 Hz), slow gamma (30-55 Hz), and high gamma (65-100 Hz). Briefly, spectrograms were constructed in a 1 s sliding 5 s window FFT of 1000 Hz LFP data for frequencies between 1-100 Hz. Power in each frequency band relative to total power was then calculated in each time window and averaged over 3-hour blocks for each animal. A slight reduction in delta power in the last 2-3 hours was observed in the ET group compared to the CT group (Fig. S3B) (F=1.741, p=0.023, group*time effect, rmANOVA). No significant effects were observed in other bands. Changing the channel to layer 5 RSCg or V1 (medial shank) did not change our results.

### Sharp-wave ripple (SWR) detection

LFPs sampled at 1000 Hz were processed by subtracting the common average reference and filtering within the ripple band (110-250 Hz). The channels with the highest variance in the ripple band were selected as the optimal SWR channel for a channel located more laterally from the midline across animals in dorsal CA1. The filtered LFP signal was then squared and smoothed with a 20 ms window to generate an RMS power signal, which was used to detect SWRs. From this power signal, peaks greater than 4 SD were detected, and then the times when the signal crossed 1.25 SD on either side of the peak were used as start and end times for candidate SWRs. Events within 20 ms were merged, and any event shorter than 20 ms was discarded. SWR amplitude was defined by the peak of the power signal, and ripple-filtered LFP peaks were used to estimate SWR frequency by averaging the peak-to-peak intervals for each event. No significant difference was seen in dCA1 SWR properties (p>0.05, unpaired t-test with Bonferroni correction) or number of SWR per minute (SWR rate) across groups (Fig. S3C; F=0.045, p=0.835, group-effect rmANOVA). SWR properties comparison on vCA1 did not show significant group differences.

### Brain state scoring

Brain state scoring was based on methods described by (*22*, *70*) and modified to fit our data. Briefly, three signals were used for scoring: theta (5-10 Hz) frequency LFP, delta (1-4 Hz) frequency LFP, and movement speed. Spectrograms were constructed using a 5 s window length and 1 s sliding window FFT of 1000 Hz LFP data for frequencies between 1 – 100 Hz. This was done on a channel in layer 5 RSCag. Periods of sleep were identified based on continuous immobility of the animal (speed < 1 cm/s) for at least 120 s. For these immobile periods, a ratio between delta and theta LFP bands was used to discriminate between SWS (NREM) and rapid-eye-movement (REM) sleep. This ratio was smoothed with a Gaussian kernel (σ=5 s), and values were scaled between 0 and 1. Periods of sleep where theta/delta ratio remained above 0.7 for at least 20 s were classified as REM; remaining sleep periods were classified as SWS. States with duration < 2 s were joined with the adjacent state to account for noise. Results were manually reviewed by the experimenter and corrected if necessary. No significant difference was observed in the duration of awake: CT: 2.56±0.12, ET: 2.6±0.11 hours (t=−0.185, p=0.85, unpaired t-test), and first 12-hours of SWS sleep: CT: 9.9±0.22, ET: 10.1±0.2 hours (t=0.69; p=0.504; Table S1). There was no significant difference in the percentage of time spent in SWS (Fig. S3D; F=0.007, p=0.935, group-effect rmANOVA) between groups.

### Functional monosynaptic connectivity analysis

We identified monosynaptic connections between neuron pairs by looking at short-latency peaks or dips (0.8-4.8 ms) of cross-correlogram (CCG), as observed in previous studies (*19*, *23*, *24*). First, we computed CCG for all simultaneously recorded neuron pairs in the range of [-50, 50] ms with a bin size of 0.2 ms. A lower frequency baseline CCG was generated by convolving the observed CCG with a partially hollow Gaussian kernel (𝜎=10 ms, hollow fraction=60%) (*71*, *72*). We estimated the probability of observed synchrony using a Poisson distribution with a continuity correction (P_fast_<0.001) (*24*). Further checks were applied to detect significant connections: (1) peak (or trough) occurred in 0.8 – 4.8ms, consistent with monosynaptic interaction; (2) peak (or trough) p-value < 0.001; (3) peak > 3 SD (or 2 SD for trough) of baseline-corrected CCG; (4) peak (or trough) width (set of contiguous bins 1 SD from baseline) should be 0.8 – 4.8ms range; (5) peak (or trough) width does not overlap with zero-lag bin (indicative of common input). A lower threshold was used for inhibitory connections because they show a broader dip, unlike sharp, transient peaks observed in excitatory connections. For excitatory connections (EE1 and EI), there is usually a peak in 0.8 – 4.8ms, whereas there is a trough in inhibitory connections (IE). Bidirectional excitatory connections (EE2) show short-latency peaks on both sides of CCG, while reciprocal E-I (EI2) have a peak and dip on either side.

For each synaptic connection passing the above criteria, synaptic strength was estimated using two methods:

1. the excess in interaction probability in 0.8 – 4.8 ms bins: 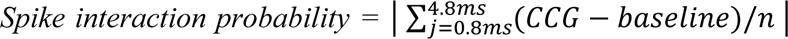, where n = number of reference neuron spikes.
2. Z-scored peak (or trough for IE) with respect to baseline CCG *z-scored peak* = | (max(CCG) – mean(baseline))/ std(baseline) | in the 0.8 – 4.8 ms. | min(CCG) | in place of max(CCG) for inhibitory connections.

For all connection types, we examine the waveform and raster plot of the detected pair to ensure this is not an overspilt cluster from spike sorting. Furthermore, common input connections between FS interneuron pairs occasionally passed the quality check and had to be sometimes manually cleaned. Total number of 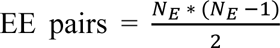 and EI pairs = 𝑁_𝐸_ ∗ 𝑁_𝐼_, where N_E_ and N_I_ correspond to the number of putative excitatory and FS interneurons in a given region x layer, respectively.

### Population vector (PV) calculation

For each animal, spikes from each good excitatory single unit were binned into 50 ms bins to generate N neurons x T time bins population vector (PV). PV was divided into 3-hour blocks, leading to 1 block for awake rest and 4 for SWS. Time bins where no neurons fired a spike (for example, down state) were excluded. Only periods with speed < 1 cm/s during awake rest and periods marked SWS based on brain state scoring were analyzed. Population sparsity, lifetime sparsity, firing rate, bursting, and PV distance analysis were done over the entire brain state epoch (awake rest and SWS) and in 3-hour blocks, roughly matching the interval between CNO injections.

### Population and Lifetime sparsity

Population sparsity (*32*, *33*) was calculated on the PV for each brain region using the following formula: Population sparsity 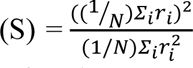; where r = mean firing rate of the *i*^th^ bin for a given neuron and N = total number of excitatory neurons. The output of the above equation is a sparsity value for each time bin, resulting in a 1 x T time bin vector.

Lifetime sparsity is a measure of the selectivity of a given neuron across its lifetime (*32*) and was calculated using the following formula: Lifetime sparsity 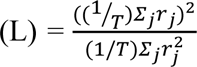; where r = mean firing rate of the *j*^th^ time bin for a given neuron and T = total time bins. PV was also used to calculate lifetime sparsity for each excitatory single unit. This results in a sparsity score for each neuron throughout the recording, resulting in a 1 x N neurons vector. Note that lower values indicate greater sparsity in both cases.

To account for differences in the number of excitatory neurons recorded across animals and brain regions, we sub-sampled individual datasets with a minimum number of neurons [n=10] for each brain region and animal 1000 times. We then averaged across these samples to allow a fair comparison of sparsity measures across animals within the same group and across groups.

### Cosine distance

To estimate the orthogonality of PVs in each region x layer combination, Cosine distance (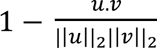, u, v are vectors,. is the dot product of u and v, ||*||_2_ is the L2-norm) was calculated across all unique time bin pairs of the PV matrix (N excitatory neurons x T timepoints) (*73*, *74*). To reduce computational demands, we applied a sub-sampling approach to calculate mean cosine distance, avoiding the computationally intensive O(N^2^) pairwise calculations for long recordings. Specifically, for each 3-hour recording epoch, pairwise distances were calculated by sub-sampling without replacement across 1000 time bins and a specified minimum number of neurons [n=10]. Distances were then averaged over 5000 random subsamples within each epoch, with each subset yielding a mean distance representing the PV orthogonality for that epoch. Note that cosine distance is insensitive to the overall scaling of activity.

### Clustering coefficient

We used graph-based analysis to determine clustering in the population activity previously described (*75*). We construct a coactivity graph between all pairs of putative excitatory cells, discarding fast movement epochs, and remove the shared influence of the general network activity on peer-to-peer coactivity using GLM. Each cell then acts as a node and the correlation between them marks the weight of the edges in the coactivity graph (N neurons x N neurons). Clustering coefficient was then computed to estimate the local coactivity structure, using the following equation: 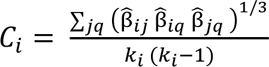, where *j* and *q* are neighbors of neuron *i*, all edge weights are normalized by the maximum edge weight in the network 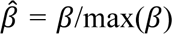, and *k_i_* is the degree of neuron *i*, which in these weighted graphs with no self-connection is equal to the number of neurons minus one (*76*). A smaller coefficient indicates that neuron i’s coactivity with its neighbors is less tightly coupled, suggesting a more decorrelated activity structure. We repeated this analysis 1000 times for each brain region and animal to determine average cluster coefficient for valid comparisons.

### Statistical analysis

Statistical analyses were performed using custom-written Python scripts or Graphpad prism software. Sample size estimation was performed using a Type I error rate (α)=0.05 and power (β)=90%, based on the mean and standard deviation estimated from a separate sample of n=4 (2 CT/2 ET) mice. Unless otherwise noted, parametric two-tailed t-test was used for unpaired data. For analysis comparing between groups across time, repeated measure analysis of variance (rmANOVA) followed by Bonferroni correction was used. Mean and standard error of mean (sem) is plotted for error bars. Central line and whiskers in the boxplot represent median, 0^th^ and 100^th^ percentile, respectively. Throughout the manuscript, statistical significance is indicated by asterisks: *p<0.05, **p<0.01, ***p<0.001. Owing to the experimental design, the experimenter was blind to the group identity.

**Fig. S1:**
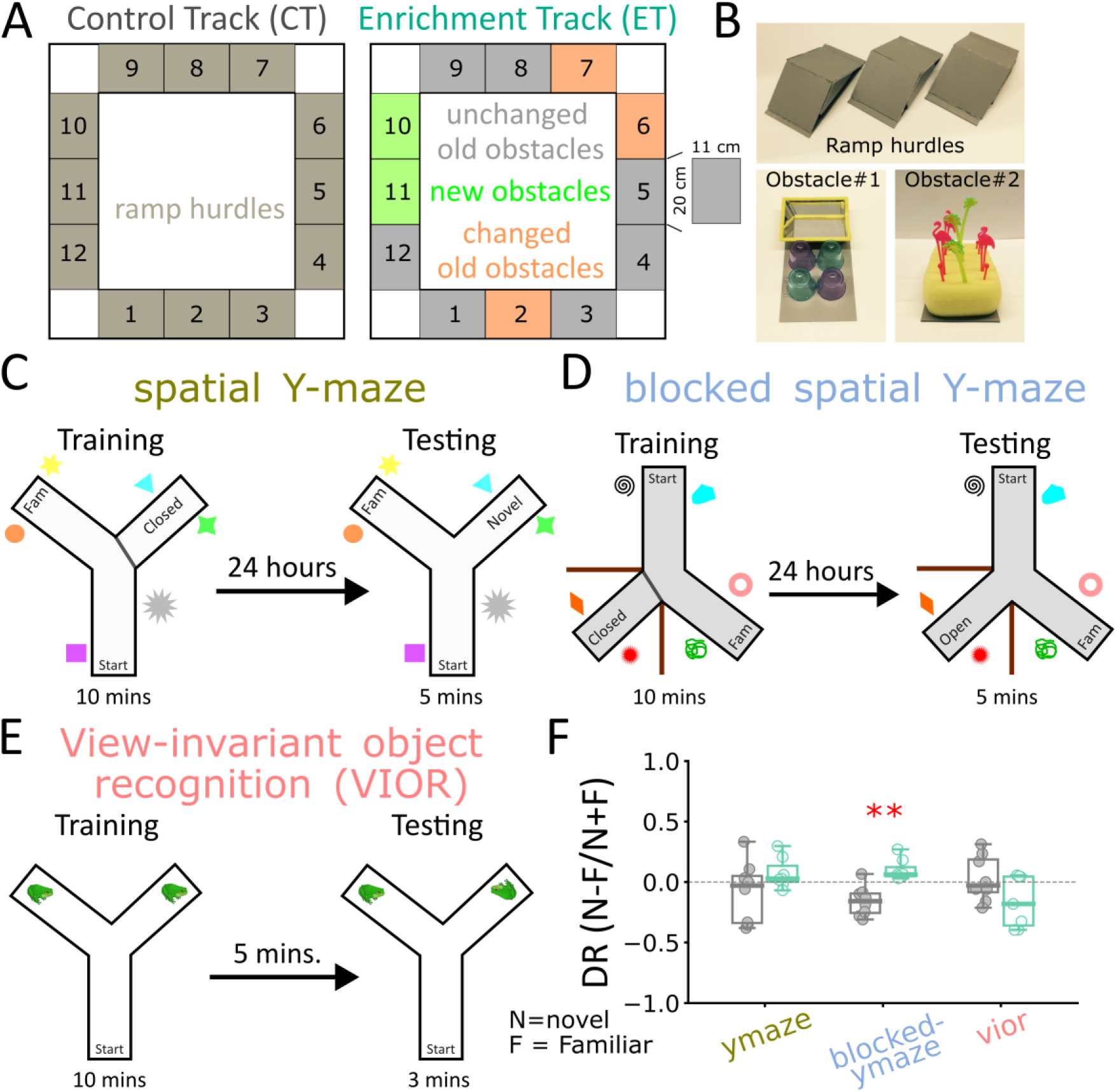
Enrichment and control track and behavior tests. (**A**) Mice ran on control (CT, left, gray) and enrichment tracks (ET, right, aquamarine). CT had 12 ramp hurdles (**B**, top), whereas ET had 12 obstacles (each 20 cm long x 11 cm wide) (**B**, bottom) through the ENR protocol. The obstacle configuration (new objects added, old obstacles rotated, translated, or swapped) was changed daily for ET mice duration. To ensure that the ENR protocol worked in our lab, n=15 (7ET/8CT) were tested in three different behavior tasks: spatial Y-maze (**C**), blocked spatial Y-maze (**D**), and view-invariant object recognition (**E**, VIOR) post-ENR to assess their spatial memory and recognition memory. Note that the animals were age-matched to experimental animals used to show changes in neural data. (**F**) Discrimination ratio 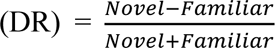, where novel and familiar indicate time spent in novel and familiar arm for each task in (C-E). ET mice performed significantly better than CT in blocked y-maze (middle, p=0.0013), suggesting better spatial novelty recognition. Dashed line indicates equal time spent in both arms.

**Fig. S2:**
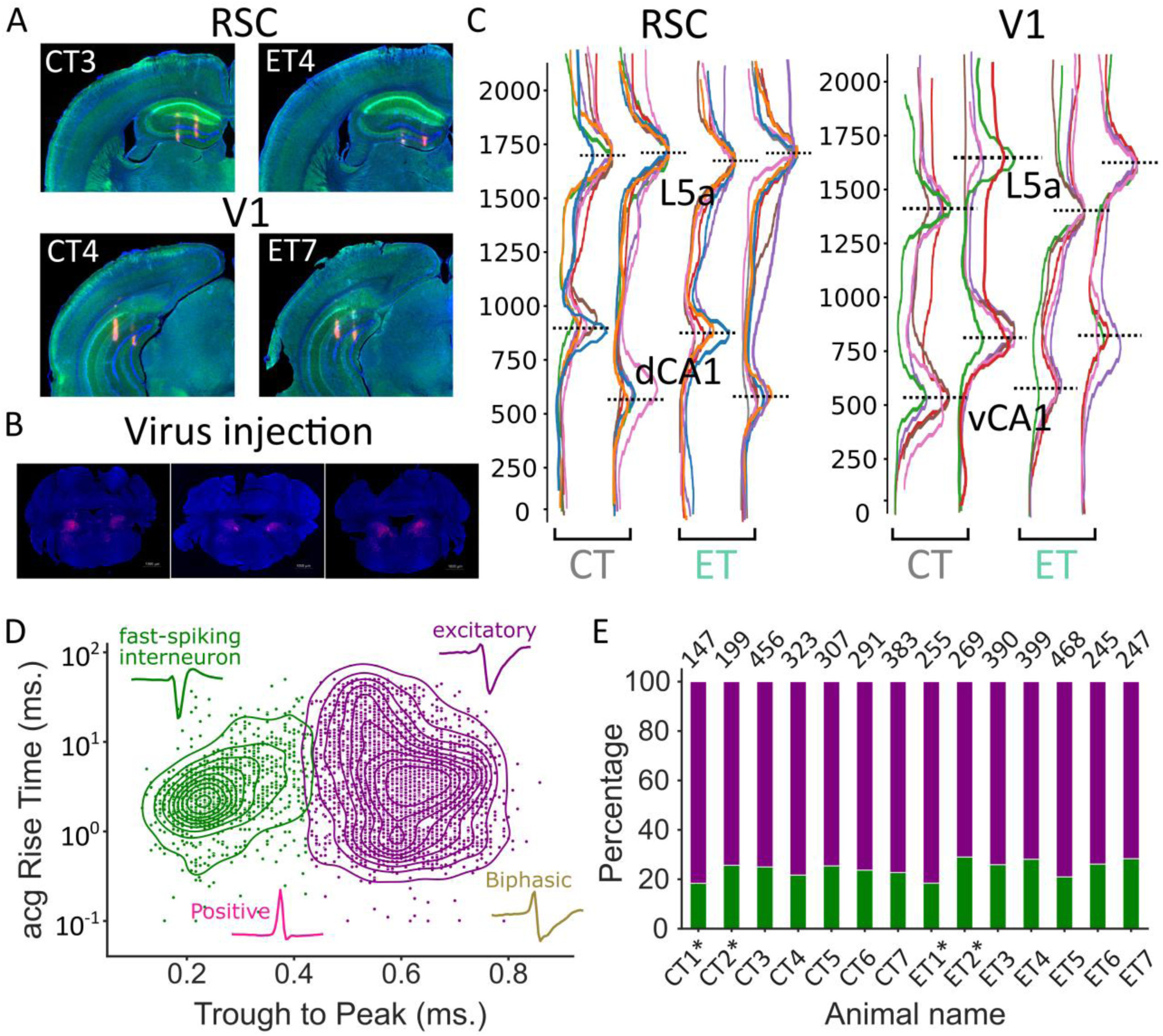
Histology, probe depth estimation using LFP, and cell-type classification. (**A**) Representative coronal sections with probe track marks (pink). Sections were stained for Wfs1 and NECAB1 to label layers 2 and 4 (both in green) in RSC (top) and V1 (bottom). (**B**) pAAV2-hSyn-DIO-hM3D(Gq)-mCherry virus expression (pink) in the parafacial zone for randomly selected coronal sections. (**C**) Relative power in the high-frequency band (100-500 Hz) for each animal’s (different colors) LFP sampled across probe depth for each shank. Notice clear peaks at layer 5a RSCg/RSCag, layer 5a V1, vCA1, and dCA1 across shanks for CT (gray) and ET (aquamarine) mice. Data for each group was affine transformed to align with a reference animal from each group. (**D**) Auto-correlogram (acg) rise time (ms) vs. mean waveform trough to peak duration (ms) (only two features shown) scatter plot showing putative FS interneuron (green) and excitatory neurons (purple). Mean waveforms of typical positive (pink), biphasic (gold), putative excitatory, and interneuron units are shown. (**E**) Percentage of cell types (only putative excitatory and FS interneurons; color same as (D)) and the total number of units (top of bar) for each animal. * indicates single probe recordings from RSC and dCA1 (CT1, CT2, ET1, ET2).

**Fig. S3:**
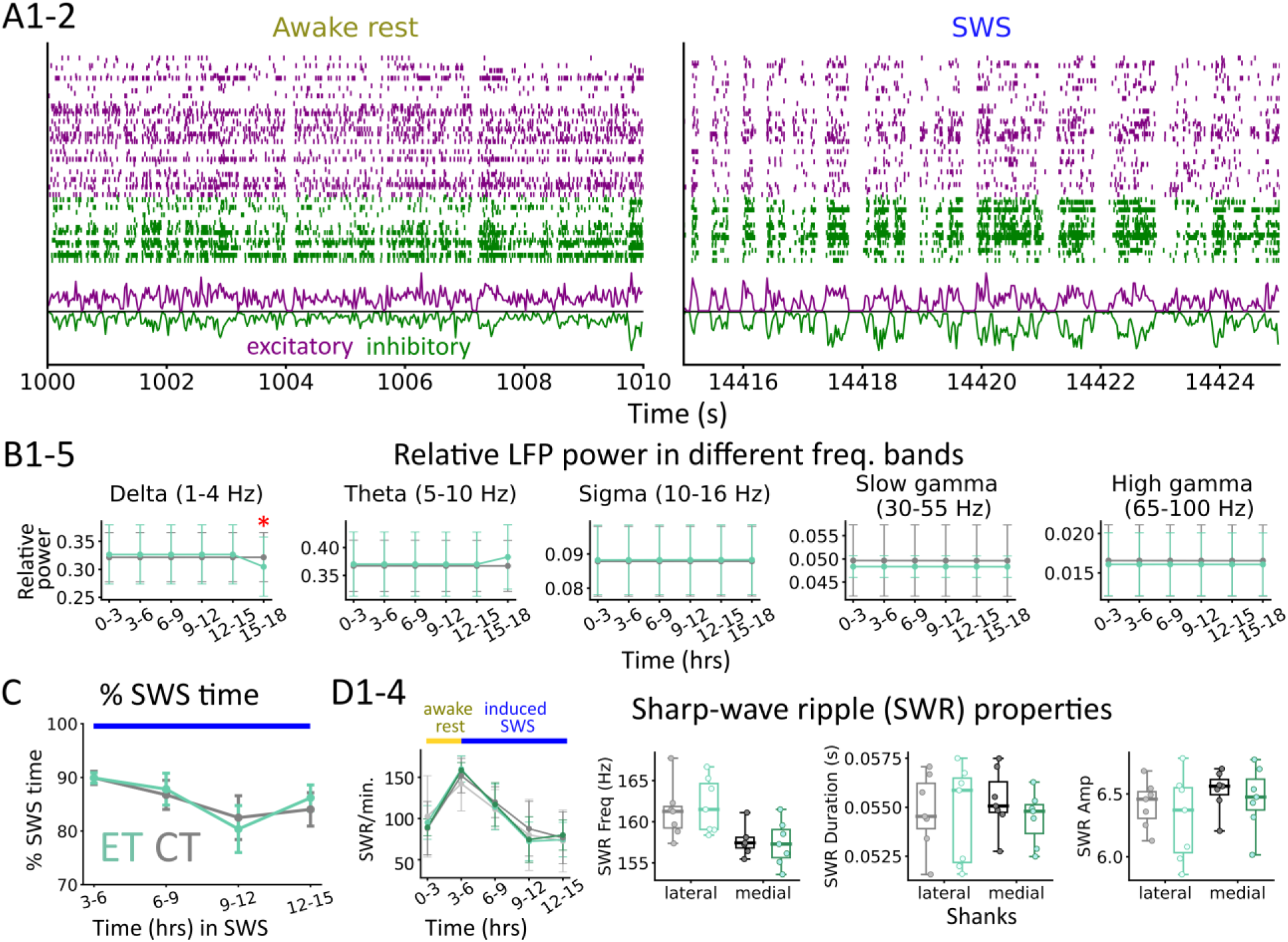
LFP parameters comparison between groups. (**A**) Raster plot showing 10 seconds of awake rest (left) and SWS (right) for excitatory (purple) and FS interneurons (green). Normalized firing activity for both neuron groups at the bottom of each subplot shows a clear E/I balance (FS interneurons activity is inverted). (**B**) Each column shows mean±sem relative power in the delta (1-4 Hz), theta (5-10 Hz), sigma (10-16 Hz), slow gamma (30-55 Hz), and high gamma (65-100 Hz), respectively for 3-hour blocks for ET (aquamarine) and CT (gray) groups. A reduction in delta power in the last 3-hour block is observed in ET mice (F=1.741, p=0.023, group*time effect, repeated-measures ANOVA), marked by red asterisk. (**C**) Percentage of time spent in SWS in the 3 to 15 hours (12 hours of induced SWS) of total recording. Each dot indicates an animal. (**D**) SWR per minute (SWR rate) for the entire recording duration, including the first 3 hours of awake rest and SWR properties: frequency (Hz), duration (s), and amplitude for channel on lateral (left) and medial (right) shank, corr. to proximal -> distal CA1. Lighter and darker hue in D1 (SWR rate) is for proximal and distal CA1, respectively. No significant group difference was seen in any variable.

**Fig. S4:**
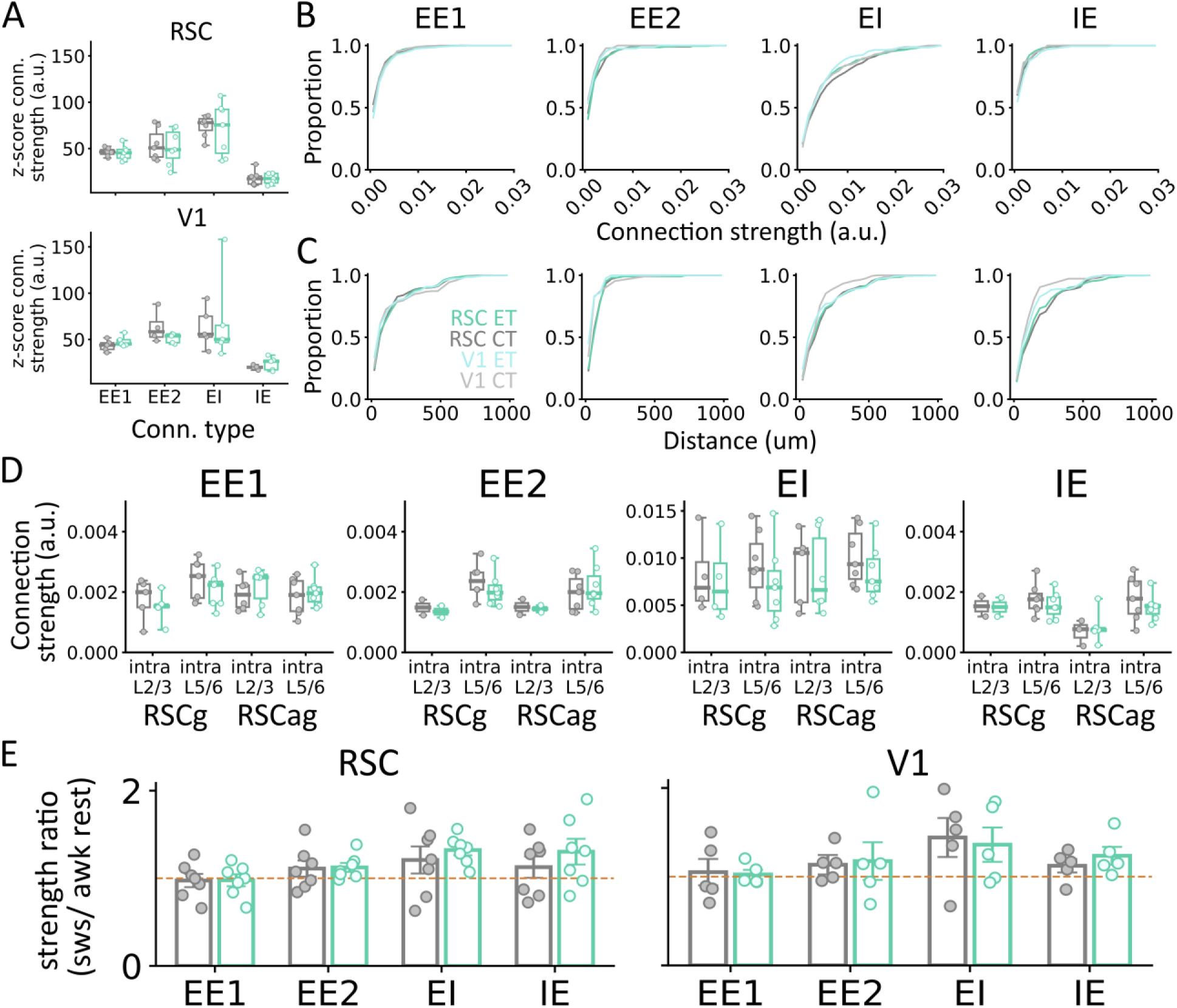
Functional synaptic connection properties between groups. (**A**) Connection strength (z-scored peak/trough with respect to baseline CCG) across connection types in RSC (top) and V1 (bottom) for ET (aquamarine) and CT (gray) groups. No significant differences were observed. Cumulative distribution of (**B**) connection strength and (**C**) distance for both groups x region (RSC ET: aquamarine, RSC CT: gray, V1 ET: turquoise, and V1 CT: silver) and connection types (EE1, EE2, EI, and IE). (**D**) Intralaminar (L2/3->L2/3, L5/6->L5/6) connection strength for EE1, EE2, EI, and IE connections in RSCg and RSCag. No significant groups differences. (**E**) Strength ratio between SWS/awake rest for EE1, EE2, EI, and IE connections in RSC (left) and V1 (right). Light brown dashed line indicates equal strength (=1). No significant group difference is observed for any connection. A small upward trend is observed from awake rest to SWS (>1) for EI, IE, and EE2 connections, but not EE1 connections.

**Fig. S5:**
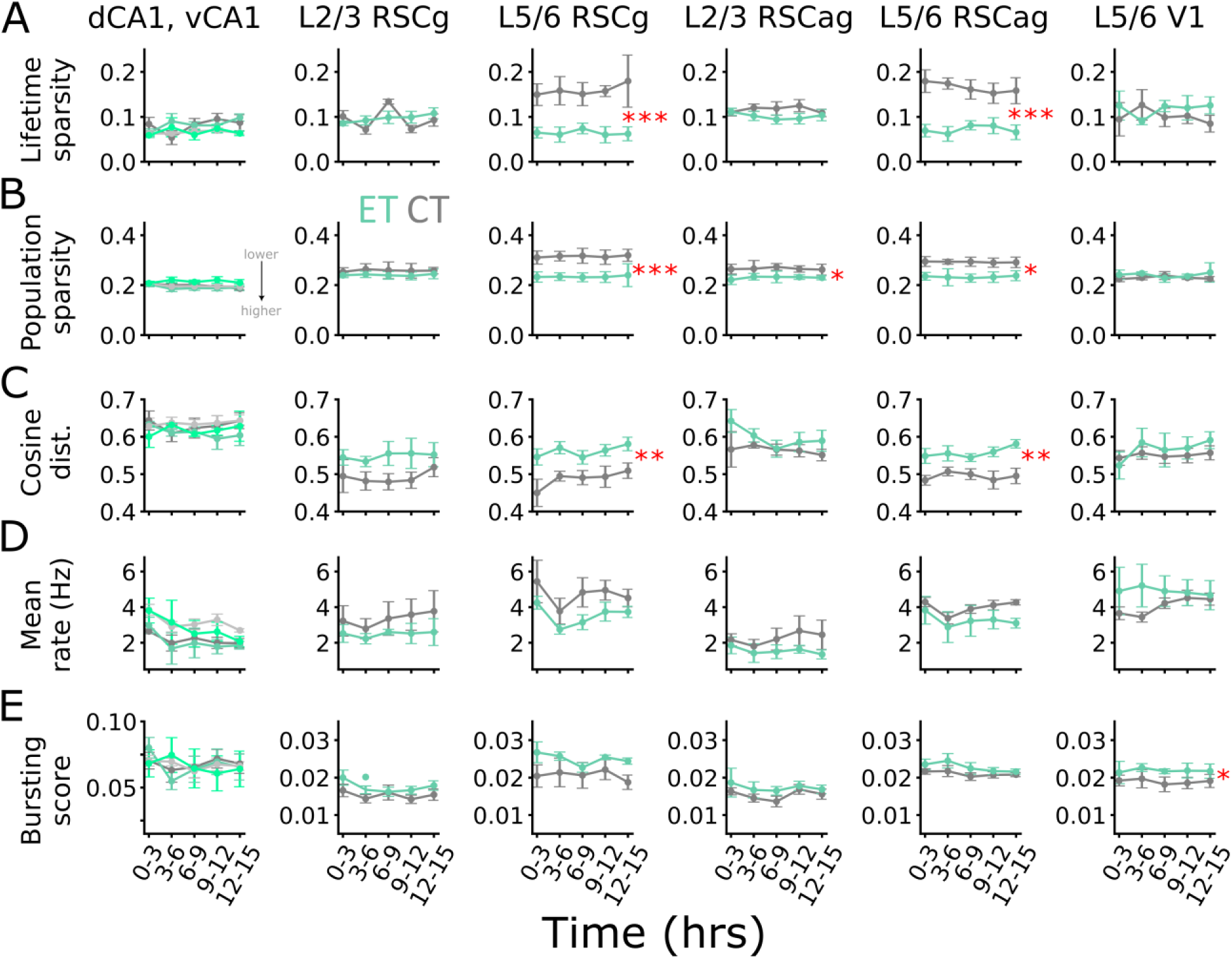
Changes in population activity dynamics across awake rest and slow-wave sleep. Mean±sem distribution for (**A**) lifetime sparsity, (**B**) population sparsity, (**C**) cosine distance, (**D**) mean firing rate (Hz), and (**E**) bursting score for excitatory neurons across brain regions: dCA1, vCA1, superficial (L2/3) and deep (L5/6) granular and agranular retrosplenial cortex (RSCg/ RSCag) and primary visual cortex (V1) for ET (aquamarine) and CT (gray) mice during awake rest: 0-3 hours and SWS: 3-15 hours (3 hours block). Lower values (closer to 0) indicate more sparsity for (A) and (B). Lighter hue is used in column 1 for both groups to indicate vCA1 data. Data from n=14 mice (7CT/ 7ET) for RSC and dCA1, and n=10 (5CT/5ET) for V1 and vCA1. Asterisks indicate group-effect for repeated-measures ANOVA, Bonferroni correction. *p<0.05, **p<0.01, ***p<0.001.

**Fig. S6:**
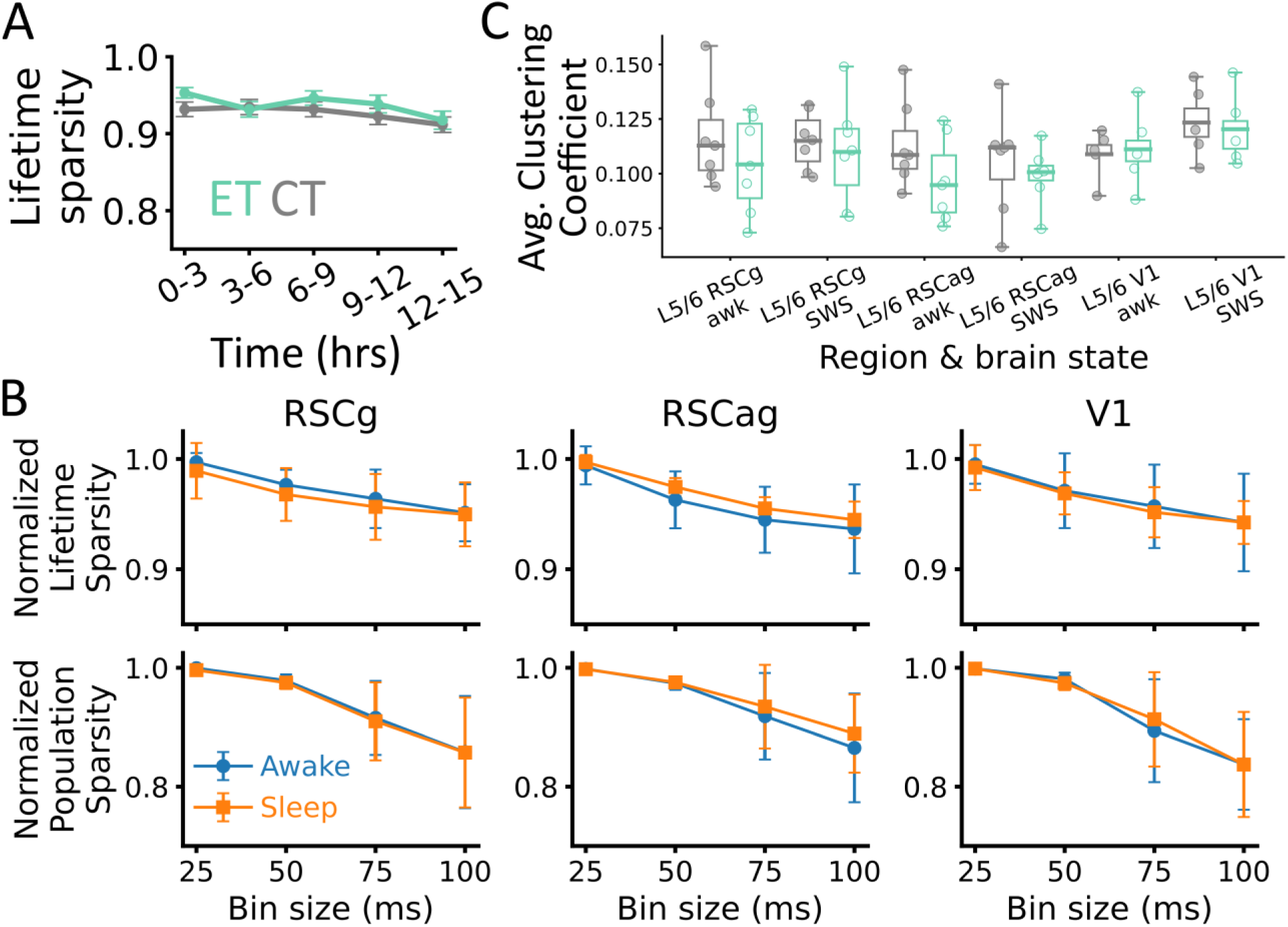
DG lifetime sparsity, neocortex sparsity vs. bin size, and clustering coefficient. (**A**) No significant difference in lifetime sparsity for dentate gyrus (DG) neurons pooled across animals for ET (aquamarine) and CT (gray) group. (**B**) Max-normalized lifetime sparsity (top) and population sparsity (bottom) for awake rest (blue) and slow-wave sleep (orange) for a range of bin size (25ms – 100ms) for generating population vectors. Values are normalized based on the maximum sparsity across all bins. (**C**) Average clustering coefficient calculated from correlation matrix for all neuron pairs across deep layers neurons for ET and CT group. Smaller value indicates that neurons do not strongly coactivate with each other.

**Fig. S7:**
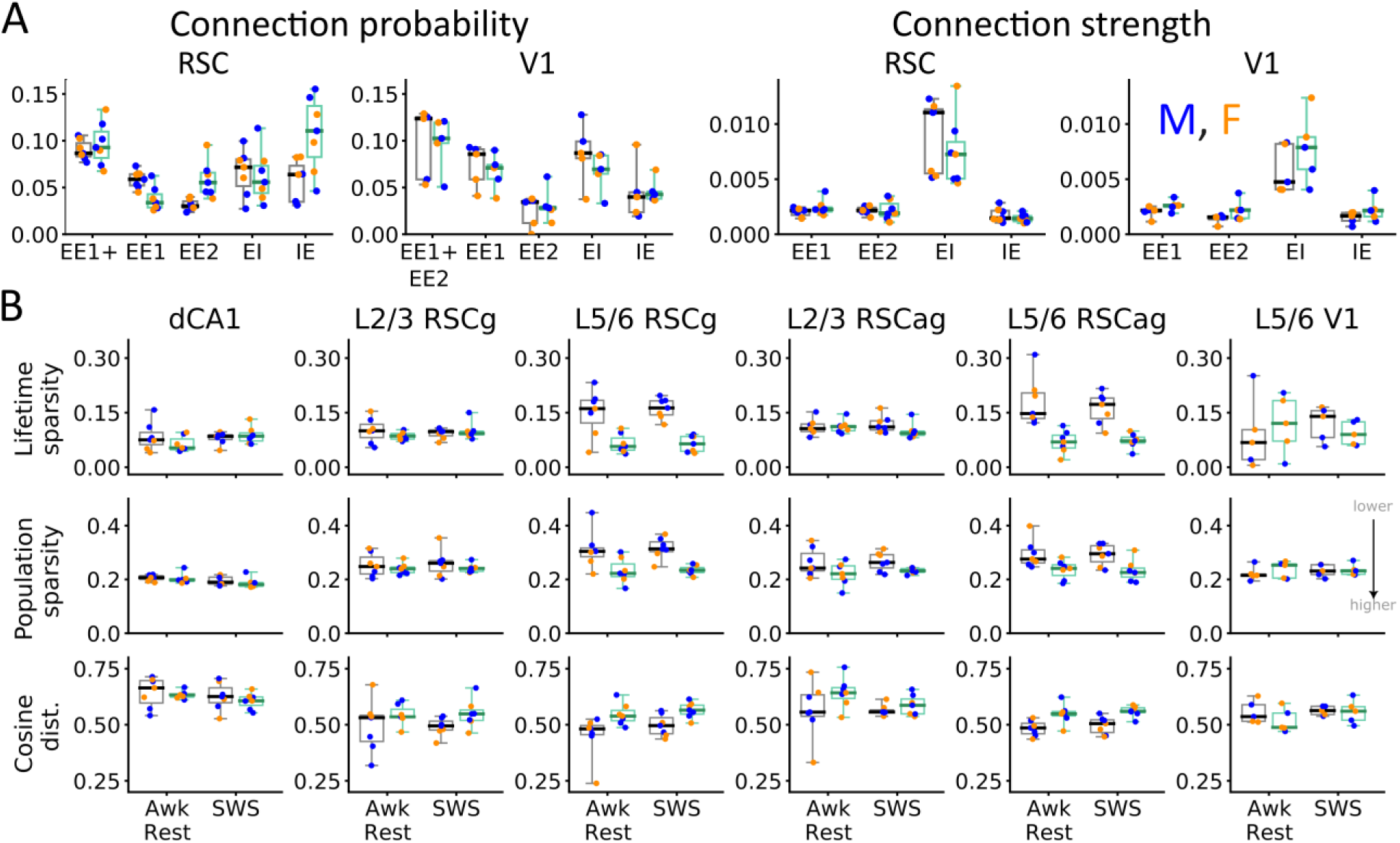
Sex comparison for connection strength and population coding dynamics. (**A**) Connection probability (left) and strength (right) for ET (aquamarine) and CT (gray). (**B**) Lifetime sparsity (top), population sparsity (middle), and cosine distance (bottom) across brain regions in ET and CT group. Each dot represents one animal, with color indicating sex (male (M)=blue and female (F)=orange). Lower values indicate more sparsity.

**Table S1:** Statistics for each animal.

| **Animal ID** | **Avg. laps** | **Total duration (hours)** | **Awake rest duration (hours) \*\*** | **SWS duration in first 12 hours \*\*** | **# single units** |
| --- | --- | --- | --- | --- | --- |
| CT1* | 47.47 | 18.09 | 2.14 | 9.15 | 147 |
| CT2* | 44.15 | 19.01 | 2.25 | 9.89 | 199 |
| CT3 | 44.3 | 22.26 | 2.75 | 9.06 | 456 |
| CT4 | 78.34 | 21.0 | 2.28 | 10.15 | 323 |
| CT5 | 91.05 | 21.84 | 2.89 | 10.47 | 307 |
| CT6 | 51.27 | 19.75 | 2.77 | 10.33 | 291 |
| CT7 | 45.34 | 23.06 | 2.88 | 10.25 | 383 |
| ET1* | 30.67 | 17.78 | 2.24 | 9.81 | 255 |
| ET2* | 27.82 | 18.51 | 2.35 | 9.2 | 269 |
| ET3 | 19.12 | 20.71 | 2.81 | 9.88 | 390 |
| ET4 | 42.2 | 23.14 | 2.33 | 10.03 | 399 |
| ET5 | 41.6 | 21.0 | 2.83 | 10.82 | 468 |
| ET6 | 34.07 | 20.58 | 2.9 | 10.44 | 245 |
| ET7 | 30.2 | 20.5 | 2.71 | 10.56 | 247 |

\* 2.5-hours awake rest compared to 3-hours in others. Also, the recording was done using a single dual-shank 256-channel probe.
\*\* Processed awake rest and SWS time after accounting for speed, power in theta band, and slow oscillations. Only the first 12-hours of data is shown for the induced sleep epoch.

**Table S2:** Functional monosynaptic properties.

| Brain region | Measured variable | EE1 |  | EE2 |  | EI |  | IE |  |
| --- | --- | --- | --- | --- | --- | --- | --- | --- | --- |
|  |  | CT | ET | CT | ET | CT | ET | CT | ET |
| RSC | Probability | 0.059 ± 0.004 | 0.038 ± 0.005 | 0.032 ± 0.002 | 0.059 ± 0.007 | 0.066 ± 0.009 | 0.062 ± 0.011 | 0.056 ± 0.009 | 0.107 ± 0.015 |
|  | Strength | 0.002 ± 0.000 | 0.002 ± 0.000 | 0.002 ± 0.000 | 0.002 ± 0.000 | 0.009 ± 0.001 | 0.007 ± 0.001 | 0.002 ± 0.000 | 0.001 ± 0.000 |
|  | Distance | 144.9 ± 10.19 | 146.9 ± 8.279 | 84.31 ± 4.372 | 75.76 ± 8.14 | 174.5 ± 12.38 | 170.91 ± 9.83 | 184.37 ± 18.34 | 172.12 ± 26.65 |
|  | Time lag | 0.002 ± 0.0 | 0.002 ± 0.0 | 0.002 ± 0.0 | 0.002 ± 0.0 | 0.002 ± 0.0 | 0.002 ± 0.0 | 0.003 ± 0.0 | 0.003 ± 0.0 |
| V1 | Probability | 0.074 ± 0.010 | 0.067 ± 0.009 | 0.024 ± 0.008 | 0.032 ± 0.008 | 0.086 ± 0.015 | 0.067 ± 0.009 | 0.045 ± 0.014 | 0.046 ± 0.006 |
|  | Strength | 0.002 ± 0.000 | 0.003 ± 0.000 | 0.001 ± 0.000 | 0.002 ± 0.000 | 0.006 ± 0.001 | 0.008 ± 0.001 | 0.002 ± 0.000 | 0.002 ± 0.000 |
|  | Distance | 131.5 ± 29.36 | 134.6 ± 12.95 | 89.03 ± 30.235 | 50.16 ± 8.451 | 126.2 ± 14.09 | 158.56 ± 22.05 | 117.45 ± 21.56 | 147.08 ± 25.93 |
|  | Time lag | 0.002 ± 0.0 | 0.002 ± 0.0 | 0.002 ± 0.0 | 0.002 ± 0.0 | 0.002 ± 0.0 | 0.002 ± 0.0 | 0.003 ± 0.0 | 0.003 ± 0.0 |

Peach color highlighted box indicates significance. Mean ± sem is listed for each measure across groups and brain states.

**Table S3:** Population coding properties.

| Region | Lifetime sparsity<br>(Awk rest, SWS) |  | Population<br>sparsity (Awk<br>rest, SWS) |  | Cosine distance<br>(Awk rest,<br>SWS) |  | Mean rate (Hz)<br>(Awk rest, SWS) |  | Bursting index<br>(Awk rest, SWS) |  |
| --- | --- | --- | --- | --- | --- | --- | --- | --- | --- | --- |
|  | CT | ET | CT | ET | CT | ET | CT | ET | CT | ET |
| **dCA1** | 0.065 ±<br>0.015,<br>0.075 ±<br>0.008 | 0.042 ±<br>0.010,<br>0.077 ±<br>0.006 | 0.207 ±<br>0.004,<br>0.187 ±<br>0.003 | 0.198 ±<br>0.010,<br>0.179 ±<br>0.005 | 0.664 ±<br>0.025,<br>0.621 ±<br>0.012 | 0.631 ±<br>0.008,<br>0.612 ±<br>0.010 | 2.657 ±<br>0.129,<br>2.031 ±<br>0.189 | 3.157 ±<br>0.453,<br>1.495 ±<br>0.303 | 0.071 ±<br>0.006,<br>0.062 ±<br>0.004 | 0.075 ±<br>0.008,<br>0.062 ±<br>0.004 |
| **vCA1** | 0.063 ±<br>0.006,<br>0.065 ±<br>0.002 | 0.060 ±<br>0.001,<br>0.061 ±<br>0.006 | 0.209 ±<br>0.004,<br>0.196 ±<br>0.001 | 0.202 ±<br>0.008,<br>0.216 ±<br>0.005 | 0.649 ±<br>0.022,<br>0.639 ±<br>0.008 | 0.605 ±<br>0.029,<br>0.626 ±<br>0.010 | 4.067 ±<br>0.305,<br>2.852 ±<br>0.134 | 4.278 ±<br>0.696,<br>2.516 ±<br>0.353 | 0.066 ±<br>0.004,<br>0.065 ±<br>0.002 | 0.075 ±<br>0.010,<br>0.056 ±<br>0.006 |
| **L2/3**<br>**RSCg** | 0.100 ±<br>0.013,<br>0.089 ±<br>0.007 | 0.084 ±<br>0.005,<br>0.091 ±<br>0.006 | 0.248 ±<br>0.017,<br>0.243 ±<br>0.012 | 0.235 ±<br>0.009,<br>0.238 ±<br>0.008 | 0.532 ±<br>0.044,<br>0.496 ±<br>0.011 | 0.535 ±<br>0.021,<br>0.539 ±<br>0.015 | 2.853 ±<br>0.854,<br>2.685 ±<br>0.435 | 2.355 ±<br>0.503,<br>2.614 ±<br>0.235 | 0.017 ±<br>0.002,<br>0.015 ±<br>0.001 | 0.018 ±<br>0.002,<br>0.017 ±<br>0.001 |
| **L5/6**<br>**RSCg** | 0.161 ±<br>0.024,<br>0.165 ±<br>0.017 | 0.053 ±<br>0.012,<br>0.059 ±<br>0.007 | 0.304 ±<br>0.026,<br>0.300 ±<br>0.012 | 0.221 ±<br>0.021,<br>0.219 ±<br>0.013 | 0.482 ±<br>0.036,<br>0.493 ±<br>0.010 | 0.533 ±<br>0.022,<br>0.567 ±<br>0.008 | 4.936 ±<br>0.710,<br>4.252 ±<br>0.324 | 4.680 ±<br>0.361,<br>3.265 ±<br>0.204 | 0.017 ±<br>0.003,<br>0.018 ±<br>0.001 | 0.024 ±<br>0.003,<br>0.025 ±<br>0.001 |
| **L2/3**<br>**RSCag** | 0.106 ±<br>0.008,<br>0.110 ±<br>0.006 | 0.108 ±<br>0.008,<br>0.094 ±<br>0.005 | 0.242 ±<br>0.020,<br>0.253 ±<br>0.010 | 0.227 ±<br>0.019,<br>0.223 ±<br>0.007 | 0.556 ±<br>0.047,<br>0.569 ±<br>0.007 | 0.647 ±<br>0.031,<br>0.585 ±<br>0.011 | 1.965 ±<br>0.341,<br>2.077 ±<br>0.331 | 1.527 ±<br>0.471,<br>1.438 ±<br>0.173 | 0.015 ±<br>0.001,<br>0.015 ±<br>0.001 | 0.017 ±<br>0.004,<br>0.017 ±<br>0.001 |
| **L5/6**<br>**RSCag** | 0.147 ±<br>0.025,<br>0.171 ±<br>0.010 | 0.069 ±<br>0.014,<br>0.072 ±<br>0.007 | 0.276 ±<br>0.020,<br>0.288 ±<br>0.008 | 0.245 ±<br>0.016,<br>0.227 ±<br>0.011 | 0.486 ±<br>0.013,<br>0.507 ±<br>0.009 | 0.553 ±<br>0.020,<br>0.565 ±<br>0.007 | 4.415 ±<br>0.262,<br>3.886 ±<br>0.136 | 3.266 ±<br>0.776,<br>2.610 ±<br>0.318 | 0.021 ±<br>0.001,<br>0.020 ±<br>0.000 | 0.023 ±<br>0.001,<br>0.021 ±<br>0.001 |
| **L5/6**<br>**V1** | 0.068 ±<br>0.044,<br>0.130 ±<br>0.014 | 0.121 ±<br>0.036,<br>0.098 ±<br>0.008 | 0.215 ±<br>0.012,<br>0.228 ±<br>0.007 | 0.253 ±<br>0.014,<br>0.233 ±<br>0.006 | 0.536 ±<br>0.023,<br>0.566 ±<br>0.010 | 0.488 ±<br>0.024,<br>0.561 ±<br>0.012 | 2.925 ±<br>0.406,<br>3.964 ±<br>0.229 | 4.907 ±<br>0.756,<br>4.630 ±<br>0.268 | 0.018 ±<br>0.002,<br>0.017 ±<br>0.001 | 0.022 ±<br>0.002,<br>0.022 ±<br>0.001 |

Awk rest = awake rest; SWS = slow-wave sleep, peach color highlighted box indicates significance. Mean ± sem is listed for each measure across groups and brain states.

